# Resolving Heterogeneity in Major Depression: Overcoupling and Undercoupling Subtypes Exhibit Differential Treatment Response and Molecular Pathways

**DOI:** 10.1101/2025.11.05.686774

**Authors:** Huaijin Gao, Yihan Ma, Rui Qian, Tao Jin, Baorong Gu, Bowen Qiu, Shaoyong Ye, the DIRECT Consortium, Manli Huang, Dan Wu, Zhiyong Zhao

**Author notes:** Corresponding to: Zhiyong Zhao, PhD, Dan Wu, PhD, Manli Huang, PhD.

## Abstract

Major depressive disorder (MDD) exhibits significant heterogeneity whose neurobiological mechanisms remain elusive. Alterations in morphological-functional coupling (MFC) have been observed in MDD. This study aims to investigate MDD subtypes based on MFC alterations and their associations with clinical symptoms and molecular basis. We identified two clinically distinct MDD subtypes in multi-center neuroimaging data (Discovery: 828 MDD/776 healthy controls; Validation: 236 MDD/86 healthy controls) by using a semi-supervised machine learning approach based on MFC changes, which were validated and robustly repeated in the validation dataset. Differences among subtypes were then examined in relation to clinical assessments, gene expression patterns, neurotransmitter and cell density, and treatment response. Subtype I (overcoupling) showed elevated MFC in high-order association cortices, linking to synaptic transmission activity and severe symptoms. Subtype II (undercoupling) demonstrated reduced MFC in primary cortices, associated with cell cycle regulation and treatment resistance. Both subtypes shared molecular signatures (cell types: astrocytes and oligodendrocytes; neurotransmitter systems: serotonergic and GABAergic receptors). Crucially, longitudinal data (33 MDD undergoing Escitalopram monotherapy) revealed that undercoupling subtype exhibited better response, implying a potential mechanism of the drug via increasing the MFC. Our work deciphers MFC-driven MDD phenotypes with distinct molecular profiles and differential treatment outcomes, suggesting potential pathways for personalized diagnosis and treatment strategies in MDD, advancing precision psychiatry.

## Introduction

Major depressive disorder (MDD) is a severe mental illness marked by significant heterogeneity across clinical symptoms, phenotypic expression, underlying etiologies, and disease progression.^1,2^ This variability complicates the identification of consistent biomarkers for diagnosis and the prediction of treatment outcomes. As a result, characterizing MDD subtypes has become a critical step toward elucidating the disorder’s underlying neurobiological mechanisms, with the potential to advance personalized, biologically-informed diagnostic and therapeutic strategies.^3^ Distinct symptom patterns in MDD may arise from divergent pathophysiological mechanisms, which limits the utility of symptom-based classifications for identifying etiologically meaningful subgroups.^1,4^ Previous neuroimaging studies have revealed MDD subtypes with different clinical outcomes or treatment responses, showing hyper- and hypo-connectivity, such as structural connectivity (SC) in the WM,^5^ or functional connectivity (FC) within the DMN.^6^ These findings may reveal the potential neurobiological substrates of key sources of heterogeneity in depression.

MDD and its subtypes have shown disrupted SC and FC in wide cortical and subcortical regions or networks, such as prefrontal lobe, sensorimotor cortex, default mode network (DMN), limbic and frontostriatal networks.^5–7^ Notably, the brain’s structural architecture and functional activity are tightly interrelated: anatomical structure constrains neural activity, while neural activity can, in turn, influence structure via plasticity and neuromodulation.^8^ Morphological-Functional Coupling (MFC)—quantifying the relationship between morphological similarity and functional connectivity^9,10^ —captures this interdependence and provides a more integrated view of brain organization. MFC has emerged as a sensitive marker for detecting subtle brain changes and underlying neuromechanism in psychiatric disorders, including MDD,^11^ and may outperform SC or FC alone in tracking disease progression and symptom expression.^12,13^ Case-control studies have reported disrupted MFC in key networks such as the DMN, attention and frontoparietal network networks in MDD, with links to clinical symptoms like suicidality and antidepressant response.^12^ However, these findings have been inconsistent across studies, possibly due to the underlying heterogeneity of MDD. Some studies reported increased MFC in sensorimotor network^14^ and decreased MFC at whole brain level^15^ in MDD compared to controls, which were not observed in other studies.^16^ To date, no study deciphered MDD subtypes based on individual-level MFC patterns.

The latest GWAS studies and transcriptomic analyses in postmortem human brains of MDD have highlighted the molecular pathways in dysregulated glutamatergic signaling and synaptic vesicular functioning, and the synaptic dysfunction-related genes like *NEGR1, DRD2, CELF4,* and *CCDC71.*^17,18^ Recent advances in whole-brain mapping of neurotransmitter systems and gene expression have unveiled unprecedented opportunities to investigate the complex relationships between neuroimaging phenotypes and underlying molecular mechanisms.^13^ Integrating multi-modal biological data, such as transcriptomic profiles and neurotransmitter distributions, with clustering algorithms and clinical symptomatology, can enhance the robustness and biological validity of data-driven subtyping approaches.^2,19^ Several studies have examined the molecular correlates of MFC alterations in MDD.^16,20^ Regional MFC alterations in MDD patients were significantly associated with the spatial distributions of neurotransmitters such as dopamine transporter (DAT) and metabotropic glutamate receptor 5 (mGluR5), as well as with gene expression patterns of *EPN2* and *MKKS.*^21^ Similarly, Long et al. reported that the neuroendocrine transporter mediated the relationship between *ABHD6* expression and MFC alterations in MDD.^16^ These findings suggest that MFC alterations may reflect complex neuropathological processes involving genetic, neurochemical, and cellular components. Recent work has leveraged genetic data^22^ to classify MDD patients into subtypes with distinct molecular signatures, such as those characterized by specific neural cell types or gene expression patterns.^23^ These distinct neuropathological pathways may underlie the variability observed in neuroimaging phenotypes.

Therefore, by integrating individual-level MFC alterations with neurotransmitter receptor distributions and gene expression profiles, we can provide convergent molecular evidence that supports the biological differentiation of MDD subtypes derived from neuroimaging data. This study aimed to identify neurobiological subtypes of MDD by examining individual-level MFC patterns. A schematic overview of the study design is presented in Figure 1. Using cross-sectional resting-state fMRI and T1-weighted imaging data from the REST-meta-MDD cohort (discovery dataset), we first applied a semi-supervised clustering approach to classify MDD patients based on their MFC alterations. We then evaluated subtype differences in demographics, clinical symptoms, MFC and the associations between MFC alterations and genetic, cell-specific transcriptomic as well as neurotransmitter profiles. Finally, we validated the subtyping results and their potential correlations with treatment-resistant depression (TRD) in an independent validation dataset, and examined treatment response to Escitalopram monotherapy in another longitudinal dataset. Our analysis revealed two distinct neurophysiological subtypes of MDD, each characterized by unique patterns of MFC alterations, along with specific associations with genetic and neurotransmitter, cellular, as well as treatment response profiles.

**Figure 1.**
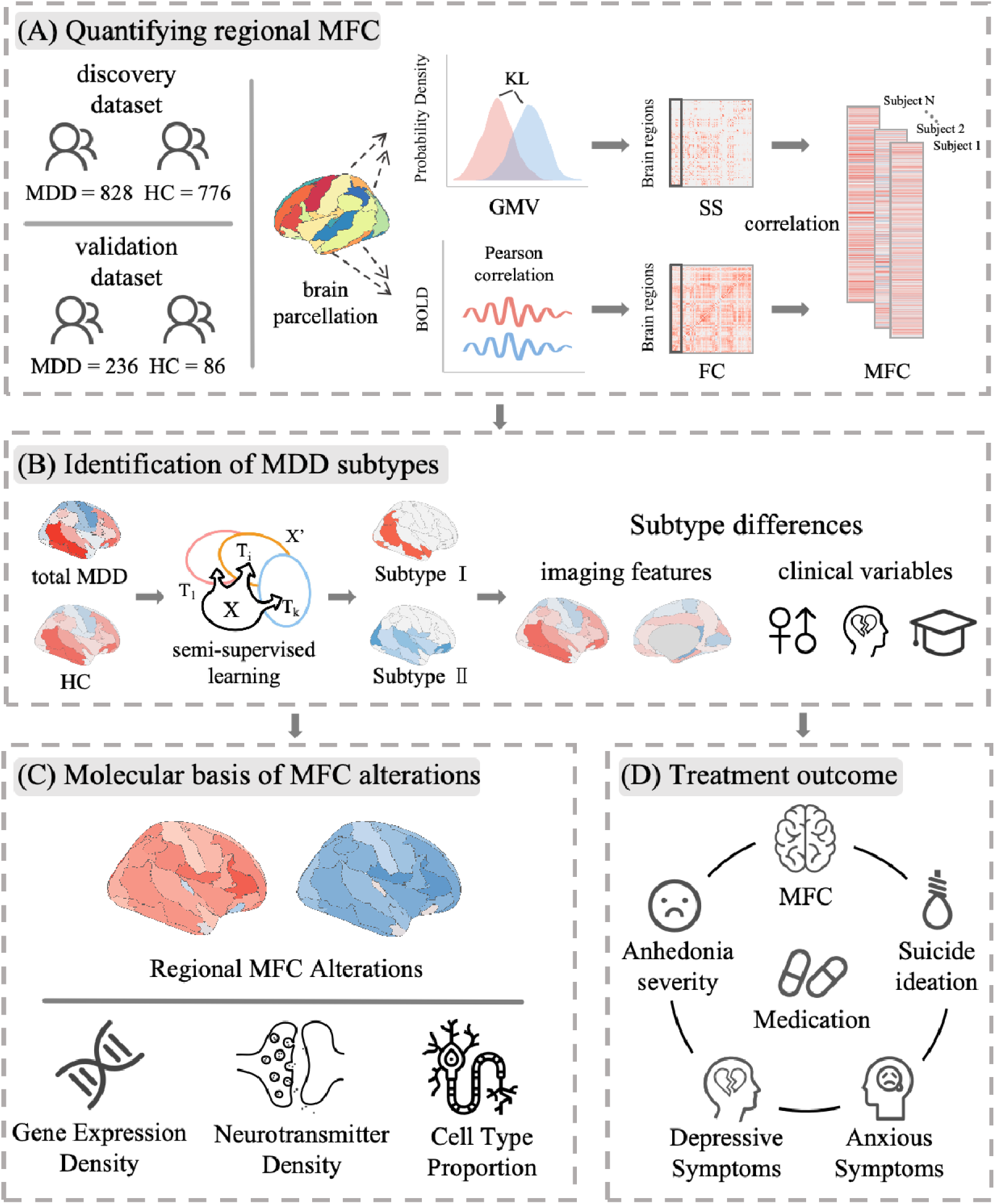
Data processing and analysis pipeline. (A) Regional MFC Quantification: morphological-functional coupling (MFC) was estimated at the individual level by calculating Spearman correlation coefficients between SS and FC across brain regions. (B) Subtype Identification and Clinical Comparison: MDD subtypes were identified based on individual-level MFC alterations. Clinical symptom differences between subtypes were subsequently assessed. (C) Molecular Characterization: To investigate the molecular basis of MFC alterations, we analyzed the associations between regional MFC and gene expression as well as neurotransmitter distribution profiles for each subtype. (D) Treatment Response Analysis: Differences in treatment outcomes were examined by comparing the responses of the two subtypes to Escitalopram monotherapy in a longitudinal dataset. Abbreviations: KL: Kullback –Leibler; MFC: morphological-functional coupling; SS: structural similarity; FC: functional connectivity; GMV: gray matter volume; HC: healthy control.

## Methods

### Participants

For the discovery dataset, resting-state functional MRI (rs-fMRI) and structural T1-weighted MRI data were obtained from the open-source REST-meta-MDD database,^24^ which includes imaging and clinical data from 1,300 MDD patients and 1,128 healthy controls (HCs). After applying quality control and exclusion criteria, the final discovery cohort consisted of 828 MDD patients and 776 HCs across 17 sites in China. Exclusion criteria were as follows: (1) missing demographic information (gender, age, or education); (2) age under 18 or over 65 years; (3) poor image quality or spatial normalization, assessed via visual inspection; (4) excessive head motion (mean framewise displacement > 0.2 mm) or insufficient brain coverage (< 90% overlap with group mask); (5) spatial correlation < 0.6 (a threshold defined as mean - 2 standard deviations) between each participant’s regional homogeneity (ReHo) map and the group mean ReHo map; and (6) sites with fewer than 10 subjects. Depressive symptoms were assessed using the 17-item Hamilton Rating Scale for Depression (HAMD-17; available for n = 728, with subitem scores available for n = 352), and anxiety symptoms were assessed with the Hamilton Anxiety Rating Scale (HAMA; n = 526). All study sites obtained approval from their local institutional review boards and ethics committees. Written informed consent was obtained from all participants.

For the validation dataset, 236 MDD patients (TRD: n = 73) and 86 HCs were recruited from The First Affiliated Hospital of Zhejiang University. Inclusion and exclusion criteria for this cohort are detailed in the Supplementary Material. A subset of 33 patients received full-dose Escitalopram monotherapy for eight weeks. MRI scans and clinical assessments—including HAMA, HAMD-24, the Beck Scale for Suicide Ideation (BSI), and the Snaith-Hamilton Pleasure Scale (SHAPS), were conducted both before and after treatment. All participants provided written informed consent following comprehensive oral and written explanations. This study protocol was approved by the local Institutional Research Board.

### Data acquisition and preprocessing

Details of the MRI scanning parameters for both the discovery (Table S16) and validation datasets are provided in the Supplementary Material. All imaging data were preprocessed using an identical pipeline implemented with the Data Processing Assistant for Resting-State fMRI (DPARSF) toolbox (http://rfmri.org/DPARSF). T1-weighted images were segmented into gray and white matter using SPM, then normalized to MNI space with the DARTEL algorithm (Ashburner, 2007). To preserve the GM volume within each voxel, the processed images were modulated using the Jacobean determinants derived from the spatial normalization by DARTEL. The resulting gray matter maps were smoothed with an 8 mm FWHM Gaussian kernel. The preprocessing of rs-fMRI data included removing the first 10 volumes, slice-timing and head-motion corrections using Friston 24-parameter model, regressing nuisance signals (WM and CSF signals), normalization, smoothing and temporal bandpass filtering (0.01–0.1 Hz).

### morphological-functional coupling calculation

MFC is computed as the correlation between FC and structural similarity, where structural similarity was derived from structural similarity networks (SSN) based on T1-weighted structural MRI, which has better accessibility, higher signal-to-noise ratio and robustness to artifacts,^25,26^ compared to that from diffusion MRI-based connectome. SSN captures local morphological features and are thought to reflect underlying cytoarchitectonic or myeloarchitectonic similarities between cortical areas at the microscale, serving as a proxy for anatomical connectivity. Here, the SSN were constructed by calculating the Kullback Leibler divergence^27^ between probability density functions of regional GMV using the Automated Anatomical Labeling (AAL) atlas, which includes 116 regions of interest (ROIs). We estimated the PDF of GMV values for each ROI based on voxel-wise GMV values from each ROI. KL divergence was then computed between the PDFs of each pair of ROIs to define the edges of the structural similarity network, and it quantifies the similarity between two probability distributions, p and q, and is defined as follows:

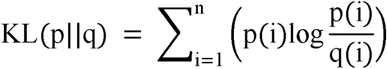

Functional connectivity (FC) networks were constructed by calculating Pearson correlation coefficients between the time series of all region pairs. The Spearman correlation coefficients for FC and structural similarity of each ROI were defined as MFC. More details are presented in Supplementary Material.

### Identifying MDD subtypes based on individual MFC alterations

Semi-supervised learning methods offer a powerful approach for identifying biologically meaningful clusters that may reflect distinct pathological trajectories, rather than simply grouping individuals based on overall data similarity, as is typical in traditional unsupervised clustering approaches. In this study, we employed a semi-supervised machine learning method called CHIMERA,^28^ which has previously been applied to classify subtypes in Alzheimer’s disease^29^ and schizophrenia,^30^ to identify MDD subtypes based on individual-level MFC, with age and gender as covariates, and site information as a grouping variable. Chimera models disease heterogeneity by mapping control and patient probabilistic distributions through multiple regularized transformations. To reduce computational demands, the input to the CHIMERA model was limited to the top 11 brain regions showing the most significant MFC differences between total MDD patients and HCs in the discovery dataset (p < 0.05, FDR-corrected), as established in prior studies.^2,6,31^ Validation analysis was conducted by applying different MFC feature selection thresholds to test the robustness of our subtyping results. Model hyperparameters included the number of clusters (K) and regularization weights (λ1 and λ2) were optimized based on two criteria: clustering reproducibility and model fit. Reproducibility was quantified using the Adjusted Rand Index (ARI),^32^ a widely accepted metric for comparing clustering consistency. Clustering experiments were conducted for K = 2, 3, and 4 clusters. For each configuration, 100 iterations of leave-10%-out cross-validation were performed. In each iteration, 90% of the MDD patients and all HCs were used to learn the transformation and form initial patient clusters. The remaining 10% of MDD patients were then assigned to clusters based on their proximity to the transformed control distribution. Based on the average ARI across the 100 iterations, the model with K = 2 yielded the most robust and reproducible clustering solution, thereby identifying two distinct MDD subtypes.

### Subtype-related demographic and clinical differences

Group comparisons between the two MDD subtypes were conducted for a series of demographic and clinical variables, including age, gender, education, disease duration, age of onset, episode status, medication status, total HAMA score, total HAMD score, and HAMD-17 item scores using two-sample t-tests for continuous variables and chi-square tests for categorical variables. Additionally, subtype differences in the association between the total HAMD score and disease duration/age of onset were examined using one-way analysis of covariance (ANCOVA) and post-hoc analyses. Furthermore, we constructed a symptom network to examine the relationships among symptom factors for each subtype. The network was based on the 17 items from the HAMD scale, and a symmetric 17×17 correlation matrix was created. Network edges were defined by correlation coefficients exceeding a fixed threshold (r ≥ 0.2), following the approach used in previous studies.^19,33^ We then analyzed the topological properties of the symptom networks for overcoupling and undercoupling subtypes.

### Subtype-related imaging differences

After identifying the MDD subtypes, we used a linear mixed model to compare MFC differences between the MDD subtypes and HC at both whole-brain and regional levels. The model describes the relationship between the response variable (here, MFC) and independent variables (here, group), while allowing for variability in the coefficients based on grouping factors (e.g., site). MATLAB’s command fitlme (https://www.mathworks.com/help/stats/fitlme.html) was utilized to test the model: y ∼1 + Diagnosis + Age + Gender + Education + Motion + (1|Site) + (Diagnosis | Site). To control for multiple comparisons, the Benjamini-Hochberg FDR correction was applied with a significance threshold of p < 0.05.

### Correlations between MFC alterations in MDD subtypes and cognition, transcriptome, neurotransmitter density as well as cell type proportion

We employed the “decoder” function to examine the relationship between the group-level t-maps, which reflect subtype-related MFC alterations compared to HCs, and the meta-analytic maps for various cognitive and neurological terms available in the Neurosynth database(https://neurosynth.org/).

To examine the relationship between MFC alterations in each MDD subtype and gene expression, we employed Partial Least Squares (PLS) regression. Specifically, we used transcriptomic data from the AHBA (http://human.brain-map.org) to link MFC changes with gene expression profiles. The transcriptomic data were preprocessed and mapped onto the 116 brain regions defined by the AAL atlas using the abagen toolbox (https://www.github.com/netneurolab/abagen). Subsequent Gene Enrichment analysis was performed on the top 1000 genes using the Metascape platform (https://metascape.org/gp/index.html#/main/step1), allowing us to identify significantly enriched biological pathways or processes associated with the MFC alterations in each subtype.

Then, Pearson’s correlations were computed between MFC alterations and the densities of 19 neurotransmitter systems from 1,200 healthy subjects, provided by JuSpace(https://github.com/juryxy/JuSpace/tree/JuSpace_v1.4/JuSpace_v1.4). The statistical significance of the correlations was determined by generating spatial permutation-based null maps, with p-values calculated from 1000 permutations and adjusted for spatial autocorrelation using partial correlation with a grey matter probability estimate.

Finally, we utilized Pearson’s correlation analyses were then conducted to examine the relationships between the spatial distributions of the six major cell types (neurons, astrocytes, oligodendrocytes, microglia, endothelial cells, and oligodendrocyte precursor cells) and alterations in MFC for each subtype. The statistical significance of these correlations was assessed using spatial permutation-based maps, with p-values obtained from 1000 permutations.

More details related to MFC-cognition, neuroimaging-transcriptome analysis, neurotransmitter information and calculation processing of cell types, are provided in the Supplementary Materials.

### Response to Escitalopram monotherapy for MDD subtypes

For clinical assessments, differences in treatment response between MDD subtypes were evaluated using a two-way repeated-measures analysis of variance (ANOVA), with time (pre-treatment vs. post-treatment) as the within-subject factor and group (Overcoupling vs. Undercoupling) as the between-subject factor. Each symptom score was analyzed as a separate dependent variable. To further investigate neurobiological alterations associated with treatment response, we examined changes in regional MFC values. For each brain region, a two-way repeated-measures ANCOVA was performed, controlling for age, gender, and education as covariates. Post hoc paired-sample t-tests were then conducted within each subtype group for significant interactions.

Moreover, for regions showing significant overcoupling or undercoupling in MDD patients (p < 0.01, FDR-corrected), we conducted one-way ANCOVA to examine MFC differences among TRD, nTRD, and HC, controlling for age, gender, and education as covariates. Then, a support vector regression (SVR) with a gaussian kernel, optimizing the regularization parameter C (range 0.1-100) and kernel coefficient γ (range 10^−3^ to 10^−1^) through grid search, was performed to classify TRD, nTRD and HC based on MFC alterations identified in the ANCOVA. Classification performance was evaluated using 5-fold cross-validation, with performance metrics including receiver operating characteristic (ROC) curves and area under the curve (AUC) values. Multiple comparisons were corrected using the Benjamini-Hochberg FDR method with a threshold of p < 0.05.

### Validation analysis

To validate our main findings, we conducted the following analyses: (1) tested the robustness of our subtyping results by applying different MFC feature selection thresholds; (2) explored the effect of medication on subtyping in unmedicated and medicated subgroups from the REST-meta-MDD dataset; (3) examined the effect of episode status on subtyping in a first-episode subgroup and a recurrent subgroup from the REST-meta-MDD dataset; (4) split the REST-meta-MDD dataset into two independent samples (N1 = 828; N2 = 776) to perform subtyping separately and assess the consistency of the identified subtypes; and (5) replicated the subtyping in the validation dataset to confirm the generalizability and reproducibility of the identified subtypes. A detailed description of the methods is provided in the Supplementary Materials.

## Results

### Demographic and Clinical Characteristics

In the discovery dataset, the MDD group had a shorter education year compared to HC group (t = −9.40, p < 0.01), but they had no significant differences in age, gender and head motion (all p > 0.05). For the validation dataset, the MDD group had significantly more females (□^2^ = 4.63, p = 0.03) and lower education levels (t = −4.87, p < 0.001) compared to the HC group. No significant differences in age were observed between the MDD and HC groups. For the treatment response exploration, 33 of the 119 MDD patients who were medicated were selected. Post-treatment analysis revealed significant reductions in HAMD (t = −10.64, p < 0.001), HAMA (t = −10.49, p < 0.001), BSI-19 (t = −4.18, p < 0.001), and SHAPS (t = −2.90, p = 0.007) scores compared to baseline. Details of the discovery and validation datasets, along with patient treatment information, are provided in the Supplemental Methods and Table S1 and Table S2.

### Differences between two subtypes in MFC alteration

Figure 2A shows the averaged MFC map across all healthy subjects. According to the optimal number of subtypes (n = 2) identified by the averaged ARI (Figure 2B), MDD was divided into two subtypes based on individual MFC alterations. The proportion of each subtype at each site is presented in Figure 2C (Subtype I: N = 388, Subtype II: N = 440). Subtype I and Subtype II exhibited structure-function overcoupling and undercoupling compared to HC at the whole-brain level, respectively (Figure 2E). Region-level analyses (Figure 2D and Table S3) further revealed that the overcoupling subtype showed increased MFC in most high-order association cortices, including the medial frontal lobe, middle temporal cortex, and angular gyrus, but undercoupling in only a few regions such as subcortical caudate and thalamus (p < 0.01, FDR corrected), while the undercoupling subtype exhibited decreased MFC in low-order primary cortices, including the precentral gyrus and inferior occipital gyrus (p < 0.01, FDR corrected). These findings demonstrate a heterogeneous pattern of MFC alterations in individuals with MDD. Additionally, when the two subtypes were mixed, the total MDD group showed less alterations than each subtype in brain regions (Figure 2D) and weaker MFC compared to HC (Figure 2E). The recurrent and first-episode drug-naïve MDD also exhibited less alterations in brain regions compared to each subtype (Figure S1 and Table S4). This further supports the necessity of MFC-based subtyping, revealing new and more MDD-related neuroimaging alteration.

**Figure 2.**
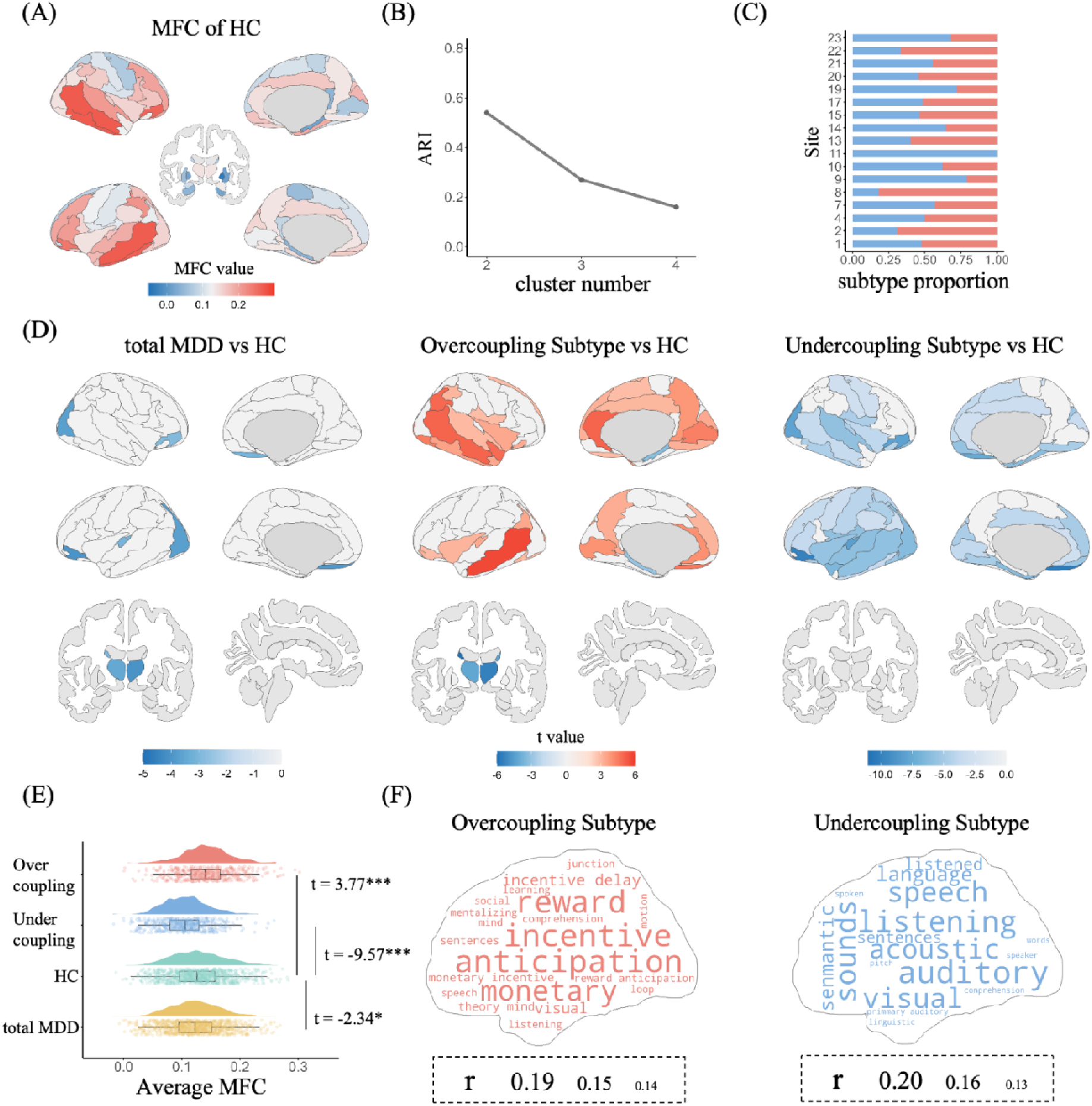
MDD subtypes and their MFC alterations compared to HC. (A) Group-level averaged MFC values in the AAL atlas for HC. (B) ARI values for all tested cluster numbers. (C) The proportion of patients in each subtype at each site. (D) MFC differences between groups at the regional level. (E) MFC differences between groups at the whole-brain level. Positive and negative t-values represent higher and lower MFC in MDD compared to HC, respectively. (F) Word clouds of the top 20 cognitive functions associated with MFC alterations. The font size of each cognitive term corresponds to the correlation coefficient (r) of the MFC alterations and the meta-analytic map generated by Neurosynth. The boxes below the word clouds indicate the correspondence between font size and the correlation coefficient. ARI: Adjusted Rand Index; AAL: Automated Anatomical Labeling. *: 0.01 ≤ p < 0.05; **: 0.001 ≤ p < 0.01; ***: p < 0.001.

Moreover, as shown in Figure 2F and Table S5, the word clouds generated by Neurosynth revealed that MFC alterations in the overcoupling subtype were primarily associated with terms related to high-order cognitive functions, such as “anticipation”, “incentive”, and “reward”, while those in the undercoupling subtype were predominantly associated with terms related to sensory and perceptual processing, such as “listening”, “visual”, and “auditory”.

### Clinical profiles differ between two subtypes

The overcoupling subtype exhibited significantly higher total HAMD scores, as well as higher scores for retardation, agitation, and hypochondriasis compared to the undercoupling subtype (Figure 3A and Table S6). Additionally, the two subtypes showed significant differences in the correlation between total HAMD scores and onset age (Figure 3B). A positive correlation was observed in the overcoupling subtype, while no significant correlation was found in the undercoupling subtype. No significant differences were observed between the two subtypes regarding illness duration. Two subtypes had no significant differences in the age, sex, education, episode (first vs. recurrent), total HAMA score, disease duration, onset age and medication status (Figure S2), suggesting the heterogeneous alterations in MDD may reflect changes in depressive symptoms rather than demographics and clinical characteristics.

**Figure 3.**
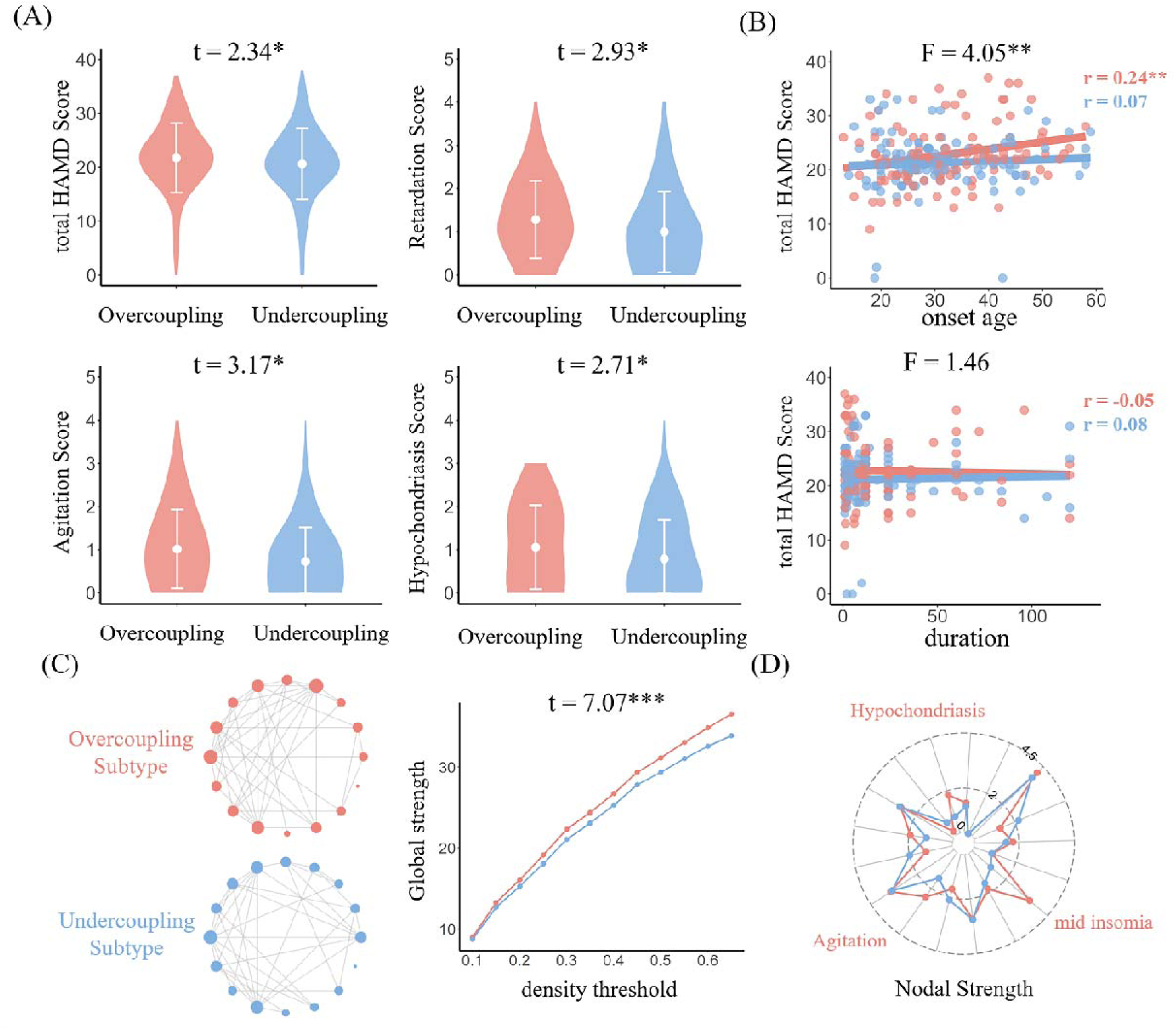
Differences between two MDD subtypes in clinical assessments. (A) Differences between subtypes in total HAMD and HAMD-17 item scores. The distributions of HAMD scores for each subtype are shown using violin plots. Each point and error bar represent the mean value and standard deviation, respectively. (B) The correlation between total HAMD score and onset age/illness duration in each subtype. The F-value indicates the magnitude of subtype differences, while the r-value represents the Pearson correlation between total HAMD score and age of onset /disease duration. (C) Symptom networks for the overcoupling and undercoupling subtype based on graph theory. (D) Radar plots show nodal strength in the symptom network for each subtype. The three HAMD-17 items with the greatest differences between subtypes are marked. HAMD: Hamilton Depression Scale. *: 0.01 ≤ p < 0.05; **: 0.001 ≤ p < 0.01.

Furthermore, using graph theory, we constructed symptom networks for two subtypes based on all 17 items of the HAMD-17. The global network strength was higher in the overcoupling subtype (global strength = 16.05) compared to the undercoupling subtype (global strength = 15.28) at a network density threshold of 0.2. This finding remained consistent across a range of network density thresholds (t = 7.07, p = 2.08×10^−5^, Figure 3C). The overcoupling subtype also showed nodal network strength differences compared to the undercoupling subtype, with the most pronounced differences observed in middle insomnia, agitation, and hypochondriasis assessments (Figure 3D and Table S7). Collectively, these findings suggest that patients with the overcoupling subtype may have more severe clinical symptoms compared to those with the undercoupling subtype.

### Transcriptomic, neurochemical and cellular signatures underlying MFC differences between MDD subtypes

The first gene component (PLS1), with the largest explained variance (overcoupling subtype: explained variance = 51.96%, p = 0.001; undercoupling subtype: explained variance = 43.87%, p = 0.02), was significantly positively correlated with MFC alterations in each subtype relative to HC (Figure 4A and B). The genes exhibiting strong correlations with MFC alterations differed between the two subtypes, with *CRTC1*, *DHRS11*, and *ZBED6* for overcoupling subtype and *SMC5*, *BTG1*, and *LOC100130* for undercoupling subtype (Figure 4C and D). Gene enrichment analysis resulted in a topologically interactive network enriched for several KEGG and GO pathways. Specifically, the overcoupling subtype’s PLS1 was enriched in biological processes related to presynapse formation, modulation of chemical synaptic transmission, and axon development (Figure 4E and Table S8). In contrast, the undercoupling subtype’s PLS1 was associated with cell cycle (morphogenesis, proliferation, division and apoptosis), and dendrite development (Figure 4F and Table S9). The network visualization effectively highlighted the functional associations and emergent patterns among the enriched terms. These findings indicate that the MFC differences between two MDD subtypes in MFC alterations may be associated with distinct gene expression profiles linked to synaptic transmission and cell cycle pathways.

**Figure 4.**
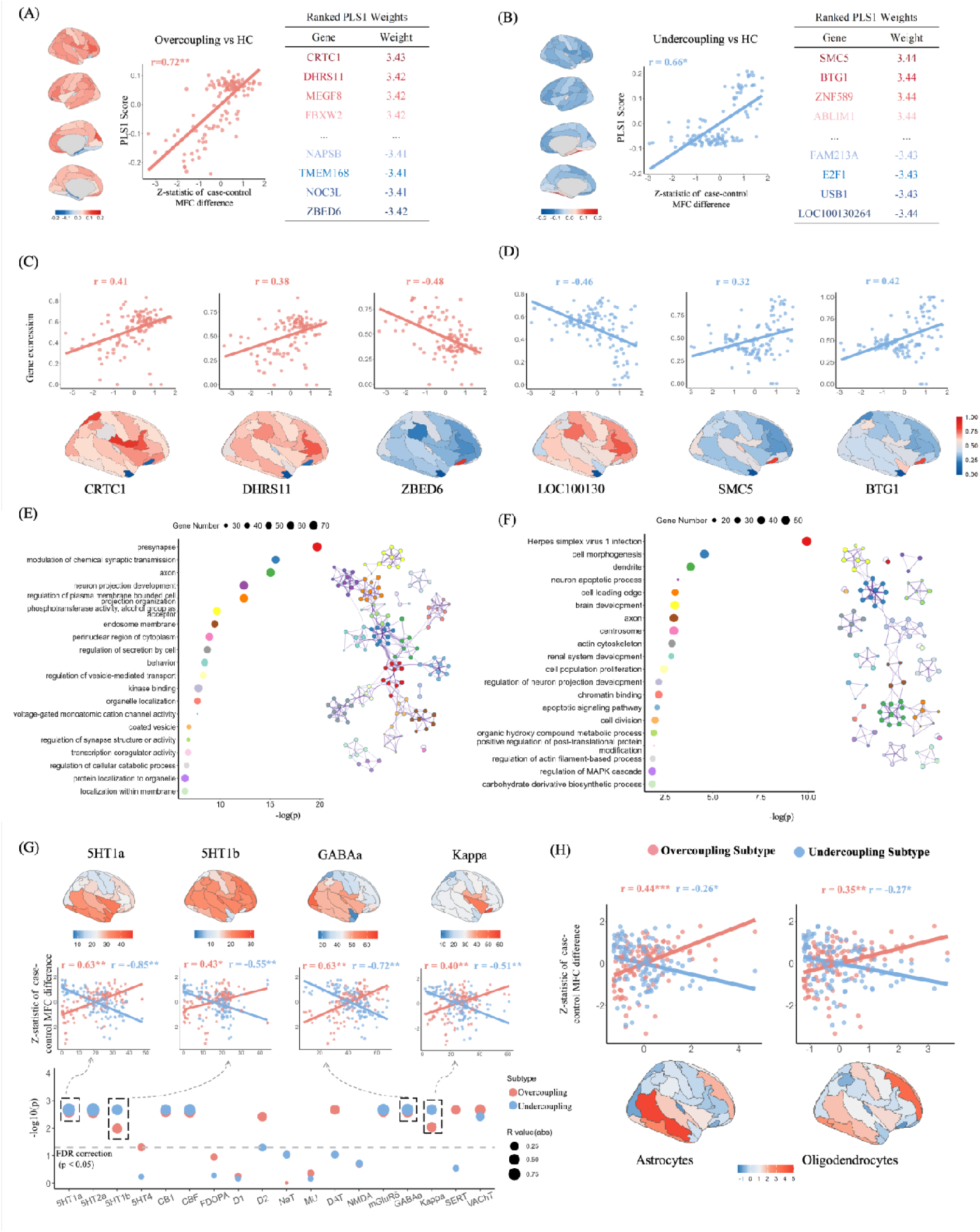
Correlations between MFC and gene expression, neurotransmitter density and cell type distribution. (A) Associations between gene expression and MFC alterations in the overcoupling and undercoupling subtype (B) compared with HC, along with the positively and negatively weighted PLS1 values of genes. The PLS1 score maps for each subtype are shown using the AAL atlas, with corresponding scatter plots provided. (C) Spatial gene expression maps with the highest absolute weights from PLS1 in the overcoupling subtype, and in the undercoupling subtype (D), along with scatter plots illustrating gene expression density across brain regions. I Enriched gene terms and pathways for altered MFC in the overcoupling subtype, and in the undercoupling subtype (F). Both intra- and inter-cluster connections are shown, delineating up to 10 terms per cluster, with each cluster distinguished by a distinct color code. (G) Spatial correlations between MFC alterations in each subtype and neurotransmitter densities across brain regions. The vertical axis represents permutation-based p-values after FDR correction, while dot sizes indicate the magnitude of the Pearson correlation coefficient. The brain maps represent distributions of four neurotransmitters showing common associations with MFC alterations in the two subtypes. (G) Spatial distributions of two cellular types and their correlations with MFC alterations for both two types. The horizontal axis represents the regional cell type distribution after z-transformation. *: 0.01 ≤ p < 0.05; **: 0.001 ≤ p < 0.01; ***: p < 0.001.

We identified significant associations between MFC alterations and neurotransmitter distributions in the two subtypes (Figure 4G and Table S10). Specifically, the MFC alterations in two subtypes shared associations with neurotransmitter systems in the serotonergic system (i.e., 5HT1a, 5HT1b, and 5HT2a), GABAa, Kappa, VaCht, mGluR5, CB1, and CSF. The opposite correlations may be derived from the overcoupling and undercoupling patterns. Furthermore, the MFC alterations in the overcoupling subtype were significantly correlated with four neurotransmitter profiles, including 5HT4, D2R, DAT, and SERT, whereas these associations were not observed in the undercoupling subtype, suggesting that dopamine systems and serotonin transporter may play key roles in underlying neurochemical mechanism of morphological-functional overcoupling in MDD.

Both astrocytes and oligodendrocytes were significantly correlated with MFC alterations in two subtypes (Figure 4H and Table S11). Moreover, the MFC alterations in the overcoupling subtype were positively correlated with the distribution of neurons (r = 0.31, p = 0.003), endothelial cells (r = 0.25, p = 0.02), and microglia (r = 0.49, p < 0.001), which were not observed in the undercoupling subtype. These findings suggest that astrocytes and oligodendrocytes are associated with MFC changes in MDD, while neurons, microglia, and endothelial cells may be specifically linked to the overcoupling.

### Differences between MDD subtypes in response to Escitalopram monotherapy

A significant group × time interaction effect was observed for the HAMD score. Post hoc paired t-tests revealed significant decreases in the total HAMD score following treatment in both overcoupling and undercoupling subtypes, with a larger change in later group. The SHAPS score showed a significant decrease after treatment in the undercoupling subtype but not in the overcoupling subtype, although it did not exhibit the significant interaction effect. Both HAMA and BSI scores showed no significant interaction effects, but they had significant decreases after treatment in two subtypes (Figure 5A). These findings demonstrate that the two MDD subtypes exhibit differential responses to escitalopram monotherapy, with a better response in the undercoupling than overcoupling subtype for depressive and anhedonia symptoms, while both subtypes show similar responses for anxious and suicidal symptoms. Significant group × time interaction effects were observed for regional MFC values in six brain regions: left orbital part of the inferior frontal gyrus, right orbital part of the inferior frontal gyrus, left olfactory cortex, right olfactory cortex, right middle occipital gyrus, and right pallidum. However, these interactions did not survive after FDR correction (p < 0.05). These interaction effects potentially support the different response to escitalopram monotherapy for two subtypes. Post hoc tests showed no significant treatment-related MFC changes within either subtype (Figure 5B and 5C).

**Figure 5.**
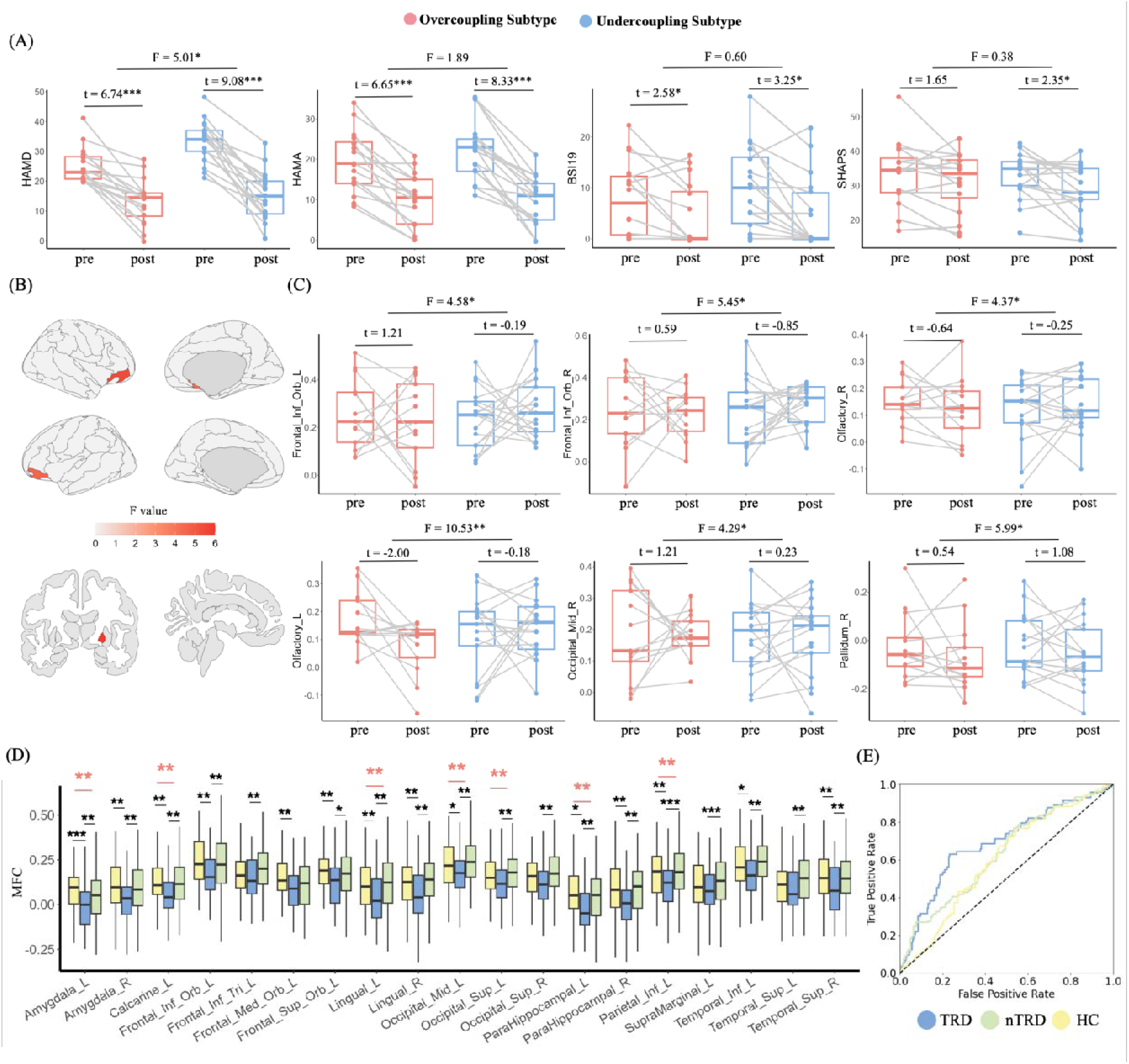
Different response to Escitalopram monotherapy across MDD subtypes. (A) Changes in clinical assessments before and after treatment for patients classified into two MDD subtypes. HAMD: Hamilton Depression Rating Scale; HAMA: Hamilton Anxiety Rating Scale; BSI: Beck Scale for Suicide Ideation; SHAPS: Snaith–Hamilton Pleasure Scale. (B) Brain regions showing significant group × time interaction effects (p < 0.05, uncorrected). (C) Changes in regional MFC values before and after treatment for patients in each subtype. Grey lines represent within-subject symptom changes over the 8-week Escitalopram treatment period. (D) Brain regions showing significant MFC differences among TRD, nTRD and HC (p < 0.05, FDR-corrected). Red asterisks denote significant F-values from ANCOVA, and black asterisks indicate significant pairwise differences. (E) Receiver operating characteristic (ROC) curves demonstrating discriminative power of MFC for distinguishing TRD, nTRD, and HC groups. Dashed line indicates chance-level performance (AUC = 0.5). nTRD: non-TRD; HC: healthy control. *: 0.01 ≤ p < 0.05; **: 0.001 ≤ p < 0.01; ***: p < 0.001.

Moreover, significant differences were observed among TRD, non-TRD (nTRD), and HC groups in the regions showing altered MFC in either overcoupling or undercoupling subtypes compared with HC (Table S12 and S13). Specifically, TRD group exhibited decreased MFC in the amygdala, calcarine cortex, lingual gyrus, occipital gyrus, parahippocampal gyrus, and parietal gyrus compared to both nTRD and HC groups, while no significant differences were observed between nTRD and HC groups (Figure 5D). Our classification analysis further confirmed that these MFC features demonstrated a superior discriminative power for TRD (AUC = 0.71) compared to the differentiation of nTRD (AUC = 0.66) and HC (AUC = 0.63) (Figure 5E).

### Validation Results

The two subtypes showed strong reproducibility across several validation analyses, underscoring the robustness of the classification. Specifically, two subtypes were consistently identified (1) in both first-episode MDD patients and recurrent MDD patients (Figure S3A-B); (2) in both reduced and enhanced feature sets with different numbers of ROIs (Figure S3C-D); (3) in two independent subsets of the REST-meta-MDD dataset (Figure S3E-F and Table S14); (4) in the unmedicated subset (Figure S3G); (5) in the validation dataset (Figure S3H and Table S15). More details related to validation are presented in Supplementary Material.

## Discussion

This study applied a data-driven, semi-supervised learning approach to identify two major subtypes of MDD based on alterations in MFC, characterized by overcoupling predominantly in high-order association cortices, and undercoupling primarily in low-order primary cortices, respectively. These subtypes differed in clinical symptom severity, treatment resistance and responses to pharmacological treatment, and their MFC alteration patterns showed distinct association with genetic profiles and shared common molecular signatures from neurotransmitter and cell types. These findings highlight the neurobiological heterogeneity of MDD and its molecular characteristics may underlie the clinical variability observed in the disorder.

### Two MDD subtypes derived from MFC alterations

Patients with overcoupling subtype exhibited increased MFC in high-order association cortices, including regions within the DMN, which is implicated in internally directed cognitive processes. In contrast, undercoupling subtype showed decreased MFC in low-order primary cortices, such as the visual and sensorimotor areas, which are essential for sensory integration and motor control. In healthy individuals, low-order primary cortices typically demonstrate stronger MFC than high-order association areas,^9,34^ reflecting their role in rapid and efficient sensory-motor processing. The elevated MFC observed in overcoupling subtype may indicate a pathological reduction in functional diversity and cognitive flexibility within the DMN. This could impair the network’s capacity to generate adaptive to external stimuli,^8,35^ potentially contributing to maladaptive, internally focused thought patterns characteristic of MDD. Conversely, the reduced MFC seen in undercoupling subtype may compromise the efficiency of sensory input processing and the relay of information along the cortical hierarchy,^8^ which could lead to deficits in basic perceptual and motor functions. Supporting these interpretations, our cognitive association analyses revealed that morphological-functional overcoupling in MDD was related to high-order cognitive functions, such as reward processing and anticipation. In contrast, the undercoupling was linked to primary perceptual functions, including vision and auditory perception. These findings align with previous reports that regional differences in MFC can robustly predict individual cognitive performance^9,35^ and are functional specificity.^8^ Collectively, these results highlight MFC as a key neuropathological marker of MDD, and distinct patterns in region-specific MFC abnormalities across subtypes may underlie the heterogeneous clinical symptoms observed in MDD and help reconcile inconsistent results reported in earlier studies.

Consistent with our subtyping findings, previous studies parsed the heterogeneity of depression using either SC or FC have also identified two MDD subtypes. These subtypes, differentiated by increased or decreased FC within the DMN, exhibited differences in depression and anxiety severity, recurrence rates, and illness duration.^36,37^ Under a normative modeling framework, two FC deviation-based subtypes—characterized by either severe positive or moderate negative deviations—showed significant differences in age, suicide and mood symptoms, and onset age.^38^ A recent study also reported opposing multilevel functional deviations across two subtypes, although no differences in symptom severity were observed.^39^ Another study identified two MDD subtypes: one showing abnormal hyperconnectivity within the ventral attention network and associated with sleep maintenance insomnia, and the other showing hypoconnectivity in subcortical and dorsal attention networks, associated with prominent anhedonia.^40^ Furthermore, SC-derived subtypes with structural network abnormalities exhibited different age of onset, symptom severity and cognitive deficits, and those with white-matter abnormalities had more neurocognitive deficit.^41^ Together, these findings support the existence of neurophysiological subtypes of MDD and provide a more objective understanding of the biological mechanisms underlying its clinical heterogeneity. Our study extends this literature by showing, for the first time, that in addition to SC and FC, MFC alterations also define two heterogeneous MDD subtypes with distinct spatial distributions.

Our network analysis demonstrated that subtyping also significantly linked to clinical symptoms, particularly in domains such as psychomotor retardation, agitation, and hypochondriasis. Overcoupling subtype had denser symptom networks, and was characterized by more severe depressive symptoms than undercoupling subtype, aligning with prior researches.^42,43^ Psychomotor disturbance (PmD), a core dimension of MDD encompassing both retardation and agitation, is known to persist even in remitted patients and is linked to greater symptom severity and poorer treatment response to antidepressants. Our findings implicate overcoupling within the DMN as a potential neurobiological correlate of PmD. This is consistent with previous studies that identified dysfunctional modulation of the SMN by the DMN as a transdiagnostic mechanism contributing to psychomotor symptoms across multiple psychiatric disorders.^44,45^ The distinct hierarchy-specific MFC alterations observed in our subtypes help explain variability in PmD presentation and antidepressant treatment response.^14^ Supporting this interpretation, overcoupling subtype demonstrated a poorer response to treatment in both total HAMD and HAMA scores compared to undercoupling subtype. In addition, prior work has implicated abnormalities in GMV and FC within the DMN regions in hypochondriasis, which have been interpreted as reflecting impaired self-referential processing,^6^ and may involve heightened unconscious emotional reactivity and increased cognitive elaboration.^46^ Our findings extend this literature by suggesting that elevated MFC within the DMN may contribute to hypochondriacal symptoms in MDD,^6,46,47^ further highlighting the role of DMN dysregulation in overcoupling subtype. Finally, we identified a significant positive correlation between age of onset and total HAMD scores in this subtype, but not in undercoupling subtype. This subtype-specific association may help reconcile inconsistencies in prior studies regarding the relationship between age of onset and depression severity.

### Molecular basis of MFC alterations in MDD two subtypes

Gene expression analysis revealed that genes associated with overcoupling were enriched in biological processes related to the presynapse and chemical synaptic transmission, while those linked to undercoupling were enriched in pathways related to HSV-1 infection and cell cycle. This aligns with previous studies linking synaptic transmission to FC abnormalities in MDD^48^ and reporting synaptic dysfunction as a core pathological feature in depressive disorders.^49^ Several studies have suggested an association between HSV recurrence and the severity of MDD symptoms, and between cell cycle regulation and antidepressant treatment.^50^ Chronic stress^51^ and FC dysfunction^52^ both implicated in disrupted cell morphogenesis — may also contribute to MDD pathophysiology. Interestingly, we identified distinct genes most strongly associated with MFC alterations in each subtype. In overcoupling subtype, *CRTC1*, a crucial gene for synaptic plasticity and long-term memory formation,^53^ had the highest weight in the PLS model. This gene has been linked to stress-induced depression-like behaviors and can directly induce such behaviors in animal models.^54^ In undercoupling subtype, *SMC5*, a gene involved in DNA repair, was most strongly associated with MFC alterations. Prior studies reported upregulation of *SMC5* in the brains of depressive-like mice, suggesting a role in alternative telomere lengthening and an inverse relationship between telomere length and MDD.^55^ Therefore, our findings suggest that distinct genetic profiles may underlie the macroscale MFC alterations observed in two MDD subtypes, offering potential targets for subtype-specific interventions.

Neurotransmitter analysis revealed that MFC alterations in two subtypes were commonly associated with multiple neurotransmitter systems, including serotonergic receptors, GABAa, Kappa, VaCht, mGluR5, CB1, and CSF. This aligns with the research connecting heightened serotonergic responsivity to agitated forms of depression.^56^ Dysfunction in GABAa receptors such as reduced expression or altered subunit composition—has been implicated in MDD pathophysiology, with some antidepressants restoring GABAergic inhibition.^57^ Similarly, reduced mGluR5 availability in MDD and its reversal following treatment, particularly in prefrontal regions, underscore its involvement in depressive symptoms and recovery.^58^ Moreover, cell type-specific analyses showed that both astrocytes and oligodendrocytes—the principal macroglial cell types—were significantly associated with MFC alterations in both subtypes. Astrocytes have been implicated in MDD through several mechanisms, including the modulation of neuroinflammation, disruption of metabolic homeostasis, and induction of neuronal hyperactivity via potassium channel dysfunction.^59^ Oligodendrocyte lineage cells, essential for myelination and axonal support, have shown functional impairments and reduced expression of myelin-related genes in individuals with depression.^60^ These observations suggest that, despite distinct neuroimaging and clinical features, overcoupling and undercoupling may share common cellular and neurochemical mechanisms contributing to their MFC alterations, underscoring the complex, multi-level pathophysiology of MDD.

### Similar and different response to Escitalopram monotherapy for MDD two subtypes

Escitalopram, a selective serotonin reuptake inhibitor (SSRI), is widely used as a first-line monotherapy for MDD. Previous studies have shown that SSRI treatment in MDD is associated with increased GABA concentrations, suggesting that modulation of GABAergic signaling may be a key neurochemical mechanism underlying treatment response.^61^ A recent genome-wide association study further indicated that genes associated with Escitalopram response in MDD were enriched in pathways related to synaptic plasticity and nervous system development.^62^ Our analyses of transcriptomic and neurochemical signatures of MFC alterations across MDD subtypes revealed shared associations with nervous system development (e.g., neuronal projection and brain development), as well as serotonin (5HT1a, 5HT1b and 5HT2a) and GABAergic receptors. These findings suggest that both MDD subtypes may share molecular substrates that underlie similar therapeutic responses to Escitalopram, particularly for anxious and suicidal symptoms.

Importantly, we also observed subtype-specific differences in treatment response. Patients in the undercoupling subtype exhibited greater reductions in both HAMD and SHAPS scores compared to those in the overcoupling subtype, indicating a potentially greater responsive to pharmacological treatment for depression and anhedonia in individuals with decreased MFC in primary cortices. This aligns with prior evidence linking reduced MFC in areas such as the middle and superior temporal gyrus to improved SSRI treatment outcomes.^14^ Transcriptomic analyses of treatment response to the Escitalopram in depression have implicated pathways related to cell proliferation, apoptosis, and inflammatory processes.^50^ Consistently, our results showed that genes associated with the undercoupling subtype were enriched in pathways related to cell cycle regulation—including morphogenesis, proliferation, and apoptosis—findings that were not observed in the overcoupling subtype. Moreover, the MFC comparisons before and after treatment revealed significant group-by-time interaction effects in six brain regions: the bilateral orbital part of the inferior frontal gyrus, bilateral olfactory cortex, right middle occipital gyrus, and right pallidum, all of which have been implicated in depression.^63^ Collectively, these findings suggest that differential responses to Escitalopram across MDD subtypes may be driven by distinct MFC patterns (i.e., overcoupling vs. undercoupling) and associated differences in cell cycle-related gene expression.

Interestingly, TRD group exhibited significantly decreased MFC compared to both HC and nTRD groups in several brain regions, including the amygdala, calcarine cortex, lingual gyrus, occipital gyrus, parahippocampal gyrus, parietal gyrus, and frontal gyrus. This aligns with previous findings indicating altered FC in the frontal cortex, amygdala, and hippocampus,^64^ as well as reduced volumes in the amygdala and frontal areas,^65^ as characteristic features of TRD. Notably, such alterations were absent in the nTRD group, suggesting that morphological-functional undercoupling may be specifically linked to treatment resistance in depression. It is worth noting that only approximately 20% of TRD patients in our sample received Escitalopram. Thus, while the undercoupling subtype may indicate a better response to Escitalopram, this observation may not generalize to other antidepressants. Conversely, patients exhibiting an overcoupling profile—associated in our findings with neurotransmitter systems such as 5-HT4, D2, DAT, and SERT—may benefit more from medications targeting these systems.^66^

Additionally, although the discovery cohort included both first-episode and recurrent cases, replication analyses using these subsets from the REST-meta-MDD dataset revealed comparable MFC alteration patterns across the identified subtypes. This suggests that the subtype classification based on MFC alterations is not dependent on illness episode status. Notably, overcoupling subtype was not replicated in medicated patients, implying it may be more sensitive to pharmacological effects. In contrast, undercoupling subtype remained consistent in both unmedicated and medicated patients, indicating greater stability. We also tested the robustness of our subtyping by varying the number of selected input ROIs and by splitting the discovery dataset into two independent subsets for separate analyses. In both cases, the subtypes remained stable. Importantly, the subtypes identified in the independent validation dataset closely resembled those in the discovery dataset, further supporting the generalizability of our findings.

### Limitations

Several important limitations in the present study should be acknowledged. First, the sample size for the longitudinal analysis was relatively small, which may limit the generalizability of treatment-related findings. Future studies should include larger cohorts to better capture individual variability in treatment response. Second, although we validated the stability of the neurophysiological subtypes using an external dataset, the validation sample was limited in size. Further replication in larger independent datasets, particularly those encompassing diverse populations in terms of ethnicity, age, and clinical presentation, is essential to establish the robustness and broader applicability of our findings. Third, research on MFC based on structural similarity in depression remains limited. Future studies should replicate our findings and examine how our MFC approach relates to conventional estimation methods, especially those utilizing diffusion MRI. Fourth, detailed medication data were unavailable for patients in the REST-meta-MDD dataset, which limited our ability to fully control for potential confounding effects of antidepressant use. Future work should investigate how different classes, dosages, and durations of antidepressant treatment may influence MFC alterations. Finally, the neuroimaging, transcriptomic, and neurotransmitter data used in this study were not obtained from the same individuals. This non-matched, integrative approach may introduce potential biases and complicate the interpretation of cross-modal associations. Future studies incorporating multimodal data from the same participants will be crucial for more precise characterization of the biological underpinnings of depression subtypes.

## Conclusion

This study identified two distinct subtypes of MDD based on MFC alterations, characterized by overcoupling in high-order association cortices and undercoupling in primary cortices. The two subtypes exhibited distinct transcriptomic signatures and shared common neurochemical and cellular substrates, underling molecular basis of the heterogeneity of clinical symptoms and response to treatment observed in MDD subtypes. These findings reinforce the importance of individualized, biology-informed treatment approaches.

## Data and Code Availability

Data of the REST-meta-MDD project are available at : http://rfmri.org/REST-meta-MDD. Neurotransmitter receptor and transporter data can be obtained online at https://github.com/juryxy/JuSpace/tree/JuSpace_v1.5/JuSpace_v1.5/PETatlas. Human gene expression data that support the findings of this study are available in the Allen Brain Atlas (https://human.brain-map.org/static/download).

