## supplementary material for "Resolving Heterogeneity in Major Depression: Overcoupling and Undercoupling Subtypes Exhibit Differential Treatment Response and Molecular Pathways"

### Supplementary Methods

#### Participants in the validation dataset

A total of 236 patients with Major Depressive Disorder (MDD) were recruited between January 1, 2023, and June 30, 2024, from outpatient psychiatry clinics at The First Affiliated Hospital of Zhejiang University, Hangzhou, China. Among which, 73 patients are defined as treatment-resistant depression (TRD), while 163 patients are defined as non-TRD (nTRD). Additionally, 86 age- and sex-matched healthy controls (HCs) were recruited from the general population through local social media platforms.

MDD diagnoses were independently confirmed by at least two trained psychiatrists based on the criteria outlined in the Diagnostic and Statistical Manual of Mental Disorders, Fifth Edition (DSM-5). Healthy controls had no personal or family history of psychiatric disorders. Inclusion criteria for MDD patients were as follows: (1) the current depressive episode met DSM-5 criteria for MDD, representing either a first episode or a relapse without prior pharmacological treatment; (2) a total score ≥ 20 on the 24-item Hamilton Depression Rating Scale (HAMD-24); (3) right-handedness; (4) minimum educational level of junior high school; (5) age between 18 and 60 years; (6) Han Chinese ethnicity. Exclusion criteria for all participants included: (1) previous treatment with antidepressant drugs or other psychotropic medications, or alternative antidepressant therapies such as transcranial magnetic stimulation or electroconvulsive therapy; (2) comorbid or past psychiatric disorders, or a family history of mental disorders in first-degree relatives; (3) secondary psychiatric disorders induced by medication or organic factors; (4) history of significant neurological disorders (e.g., epilepsy, cerebral infarction), endocrine disorders (e.g., diabetes), or other major medical illnesses (e.g., cardiovascular disease, hypertension); (5) history of alcohol or substance abuse or dependence; (6) presence of contraindications for magnetic resonance imaging (e.g., metal implants, claustrophobia).

Depression severity was assessed using the HAMD-24, and anxiety symptoms were evaluated using the Hamilton Anxiety Rating Scale (HAMA). Suicidal ideation was measured using the Beck Scale for Suicide Ideation (BSI), while anhedonia severity was assessed with the Snaith–Hamilton Pleasure Scale (SHAPS).

**MRI Data Acquisition in the validation dataset**

All MRI scans were conducted at a GE Signa HDXT 3.0T MRI system equipped with a 32-channel head coil. High-resolution 3D T1-weighted images were acquired using a sagittal 3D Brain Volume (BRAVO) sequence with the following parameters: repetition time (TR) = 7.8 ms, echo time (TE) = 2.99 ms, inversion time (TI) = 1100 ms, flip angle = 7°, field of view (FOV) = 256 × 256 $mm^2$, matrix size = 256 × 256, slice thickness = 1 mm, number of slices = 192, and total acquisition time = 5 minutes 33 seconds.

Resting-state functional images were obtained using a 2D gradient-recalled echo (GRE) echo-planar imaging (EPI) sequence with the following parameters: TR = 2000 ms, TE = 30 ms, flip angle = 90°, FOV = 220 × 220 $mm^2$, matrix size = 64 × 64, slice thickness = 4 mm, inter-slice spacing = 0.6 mm, number of slices = 33, number of volumes = 180, and total acquisition time = 6 minutes.

**Structural-functional coupling calculation**

Using a kernel density estimator toolbox in MATLAB (https://www.mathworks.com/matlabcentral/fileexchange/14034-kernel-density-estimator), we estimated the PDF of GMV values for each ROI. The optimal bandwidth for the estimator was selected automatically, and a conservative n value was used based on prior studies[100,107]. KL divergence was then computed between the PDFs of each pair of ROIs to define the edges of the structural similarity network, and it quantifies the similarity between two probability distributions, p and q, and is defined as follows:

$$KL(p||q) = \sum_{i=1}^{n}\left(p(i)\log\frac{p(i)}{q(i)}\right)$$

Since $KL(p||q)$ is not equal to $KL(q||p)$, the similarity between the two PDFs is calculated using a symmetric KL divergence, which is defined as follows:

$$KL(p,q) = \sum_{i=1}^{n}\left(p(i)\log\frac{p(i)}{q(i)} + q(i)\log\frac{q(i)}{p(i)}\right)$$

Finally, to constrain the value to a range from 0 to 1, we employed the following transformation:

$$KLS(p,q) = e^{-KL(p,q)}$$

Thus, a 116 × 116 structural similarity matrix was obtained for each participant.

Functional connectivity (FC) networks were constructed by calculating Pearson correlation coefficients between the time series of all region pairs. To improve normality, the correlation coefficients were transformed using Fisher's r-to-z transformation. The Spearman correlation coefficients for FC and structural similarity of each ROI were defined as MFC [18,24].

**Correlations between MFC alterations in MDD subtypes and cognition**

To investigate the association between subtype-specific MFC alterations and cognitive processes, we utilized Neurosynth[1] (https://neurosynth.org/) for meta-analytic analysis. Specifically, we employed the "decoder" function to examine the relationship between the group-level t-maps, which reflect subtype-related MFC alterations compared to HCs, and the meta-analytic maps for various cognitive and neurological terms available in the Neurosynth database. The top 20 terms with the highest absolute correlation coefficients, as derived from the "decoder" function, were selected for presentation. These terms represent cognitive and neurobiological processes that are most strongly associated with the observed MFC alterations in each MDD subtype, providing insight into the underlying functional implications of these structural-functional interactions.

**Transcriptomic data preprocessing**

Transcriptional data were obtained from the Allen Human Brain Atlas (AHBA; https://human.brain-map.org/), which provides microarray-based gene expression profiles from postmortem brain tissue samples. The dataset included 3,702 samples collected from six adult donors, with expression data covering 20,737 genes and 58,692 probes. Preprocessing of the AHBA dataset followed the standardized pipeline proposed by Arnatkevic et al[2]. The procedure consisted of the following five steps: (1) Probe-to-gene annotations were verified and updated using the Re-Annotator toolkit[3]; (2) Probes with expression levels not exceeding background noise in at least 50% of the tissue samples were excluded; (3) For each gene, the probe showing the highest correlation with RNA-seq data was retained; (4) Tissue samples were coregistered to the Automated Anatomical Labeling (AAL) 116 atlas using a spatial threshold of 2 mm Euclidean distance from the nearest parcel; (5) Gene expression values were spatially normalized across samples for each donor using a sigmoid transformation.

The resulting gene expression matrix (116 brain regions × 15,633 genes) was then used for further analysis. PLS regression identifies linear combinations of weighted gene expression scores (predictor variables) that best predict the MFC alterations (response variables)[4]. The first PLS component (PLS1), which explains the maximum variance, was tested for statistical significance using 1000 permutations, incorporating surrogate maps generated by spin rotations for cortical regions and resampling for subcortical regions via brainsmash [5]. To correct for estimation errors in the gene weightings, a bootstrapping method was applied during the PLS1 analysis for each group. Two separate PLS1 gene lists were generated for Subtype 1 and Subtype 2, with genes ranked based on their corrected weights reflecting their contribution to the PLS component.

**Correlation between neurotransmitter density and MFC alterations**

Pearson's correlations were computed between MFC alterations and the densities of various neurotransmitter systems, including the serotonin 5-hydroxytryptamine receptor (5-HT), dopamine receptors (D1, D2), dopamine transporter (DAT), dopamine synthesis capacity (FDORA), gamma-aminobutyric acid (GABAa), noradrenaline transporter (NAT), metabotropic glutamate receptor 5 (mGluR5), serotonin transporter (SERT), kappa opioid receptor (Kappa), vesicular acetylcholine transporter (VAChT), cannabinoid receptor (CB1), N-methyl-D-aspartic acid receptor (NMDA), μ-opioid receptors (MU), and cerebral blood flow (CBF). The statistical significance of these correlations was determined by generating spatial permutation-based null maps, with p-values calculated from 1000 permutations and adjusted for spatial autocorrelation using partial correlation with a grey matter probability estimate.

**Correlation between cell type proportion and MFC alterations**

We utilized comprehensive whole-brain reference maps of cellular abundance for six major cell types—neurons, astrocytes, oligodendrocytes, microglia, endothelial cells, and oligodendrocyte precursor cells—reconstructed by Pak et al.[6]. Their approach employed a deconvolution method based on feature gene decomposition, which utilizes known genetic markers to quantify the proportions of these cell types. Using transcriptomic data from the AHBA, Pak et al. calculated the cell type densities across 116 brain regions defined by the AAL atlas. Deconvolution was performed using the BRETIGEA software package (https://github.com/andymckenzie/BRETIGEA)[7], enabling the generation of spatially resolved cell type distributions throughout the human brain[6].

**Validation analysis**

***Different thresholds in the MFC feature selection***

In the main analysis, we selected the MFC of 11 ROIs that showed the most significant differences between total MDD patients and healthy controls (HCs) in the REST-meta-MDD dataset ($p < 0.05$, FDR-corrected). These ROIs were used as input features to reduce computational demands. To evaluate the robustness of the subtyping results, we repeated the analysis using alternative ROI sets (9 and 13 ROIs).

The semi-supervised machine learning algorithm was re-applied to classify MDD subtypes based on MFC, with age and gender included as covariates and imaging site treated as a grouping variable. Following subtype identification, a linear mixed-effects model was used to compare regional MFC differences between subgroups. To evaluate the stability of the findings across different ROI selections, we computed Pearson correlation coefficients between the T statistics from these alternative models and those obtained in the primary analysis.

***Medication and Episode effects on results***

To assess the potential influence of medication status and episode status on MDD subtyping, we repeated the subtyping procedure and reanalyzed the heterogeneity of MDD subtypes. Specifically, we selected 298 drug-naïve patients and 121 first-episode patients who were receiving medication from the REST-meta-MDD dataset. The semi-supervised machine learning method was re-applied to classify MDD based on MFC, with age and gender included as covariates and imaging site treated as a grouping variable. To further evaluate the effect of episode status, we performed the same subtyping analysis separately on two subgroups: first-episode MDD patients (n = 400) and recurrent MDD patients (n = 208). Following subtype identification, regional MFC differences between subtypes were examined using a linear mixed-effects model. To evaluate the consistency of the findings across subgroups, we calculated Pearson correlation coefficients between the T statistics obtained from the drug-naïve, medicated, first-episode, and recurrent subgroups and those from the main analysis.

***Robustness Analysis through Half Splitting the REST-meta-MDD Dataset***

To further evaluate the robustness of our subtyping results, we repeated the subtyping procedure by dividing the REST-meta-MDD dataset into two independent subsets based on site distribution. Specifically, data from sites 1 to 15 (n = 828) constituted the first subset, while data from sites 17 to 23 (n = 776) formed the second subset. Detailed descriptions of each site are provided in Table S14. For each subset, input

ROIs were independently selected based on regions showing the most significant MFC alterations in total MDD patients compared to HC ($p < 0.05$). This process resulted in the identification of 15 ROIs for both subsets. The subtyping analysis was then performed separately within each subset. To quantify case-control differences in MFC at the regional level for the identified subtypes, T statistics were computed using a linear mixed-effects model for each subset. Consistency between the two analyses was assessed by calculating Pearson correlation coefficients between the T statistics derived from the two subsets.

***Reproducibility Analysis in the Validation Dataset***

To further assess the robustness of our subtyping results, we applied the subtyping procedure to an independent validation dataset from The First Affiliated Hospital of Zhejiang University. Due to the relatively small sample size of this dataset, identifying input ROIs solely from these data could compromise statistical power and increase variability, potentially resulting in unstable feature selection. Moreover, using a limited number of ROIs as input features in the semi-supervised learning model could reduce the reliability of subtype identification. To address these limitations, we combined ROIs identified from both the validation dataset (11 ROIs; $p < 0.05$) and the large REST-meta-MDD dataset (11 ROIs; $p < 0.05$, FDR-corrected). After removing overlapping regions, a total of 19 unique ROIs were selected as input features for subtyping. The same semi-supervised subtyping procedure was then applied to the validation dataset. To evaluate the consistency of the subtyping results, we computed Pearson correlation coefficients between the T statistics derived from the validation dataset and those obtained in the primary analysis.

### Supplementary Results

#### Validation results

The two subtypes showed strong reproducibility across several validation analyses, underscoring the robustness of the classification. Specifically, (1) two subtypes were consistently identified in both first-episode MDD patients (overcoupling subtype: r = 0.92, p < 1.00×$10^{-16}$; undercoupling subtype: r = 0.95, p < 1.00×$10^{-16}$) and recurrent MDD patients (overcoupling subtype: r = 0.82, p < 1.00×$10^{-16}$; undercoupling subtype: r = 0.92, p < 1.00×$10^{-16}$) (Figure S3A-B); (2) when different numbers of ROIs were used, the subtypes remained highly consistent with the main results. This consistency was evident in both reduced (9 ROIs: overcoupling subtype, r = 0.91, p < 1.00×$10^{-16}$; undercoupling subtype, r = 0.93, p < 1.00×$10^{-16}$) and enhanced (13 ROIs: overcoupling subtype, r = 0.88, p < 1.00×$10^{-16}$; undercoupling subtype, r = 0.93, p < 1.00×$10^{-16}$) feature sets (Figure S3C-D); (3) when the REST-meta-MDD dataset was divided into two independent subsets and subtyping was performed separately, the two subtypes were still consistent, further supporting their robustness (overcoupling subtype: r = 0.48, p = 6.05×$10^{-8}$; undercoupling subtype: r = 0.43, p = 1.70×$10^{-6}$) (Figure S3E-F and Table S14); (4) two subtypes were successfully replicated in the unmedicated subset, with high correlation values (overcoupling subtype: r = 0.92, p < 1.00×$10^{-16}$; undercoupling subtype: r = 0.94, p < 1.00×$10^{-16}$) (Figure S3G). (5) two subtypes identified in the validation dataset exhibited significant similarity to those derived from the discovery dataset, reinforcing their generalizability (overcoupling subtype: r = 0.37, p = 4.82×$10^{-5}$; undercoupling subtype: r = 0.34, p = 1.84×$10^{-4}$) (Figure S3H and Table S15).

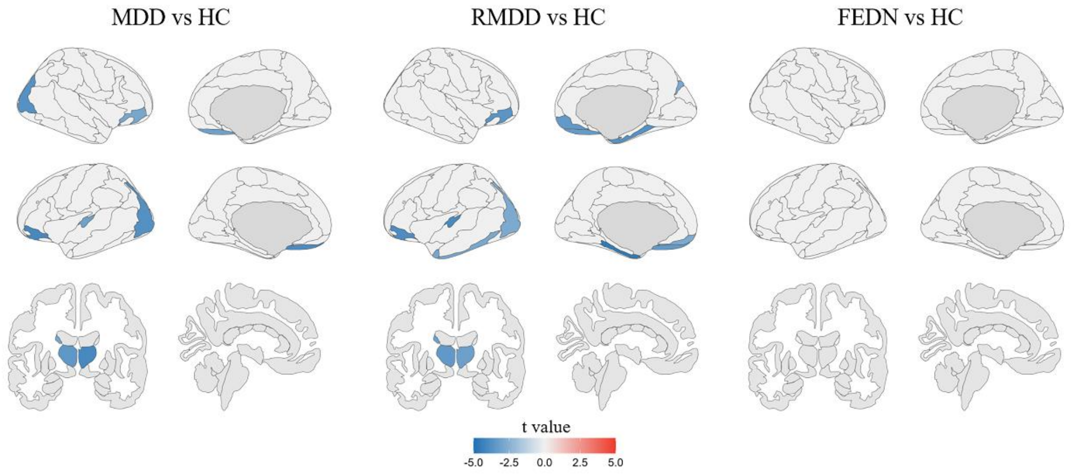


**Figure S1. Regional MFC differences between groups.** Positive and negative t-values indicate higher and lower MFC in MDD patients relative to healthy control (HC), respectively. Only regions that survived false discovery rate (FDR) correction ($p < 0.05$) are displayed. RMDD: Recurrent major depressive disorder; FEDN: First-episode, drug-naïve patients.

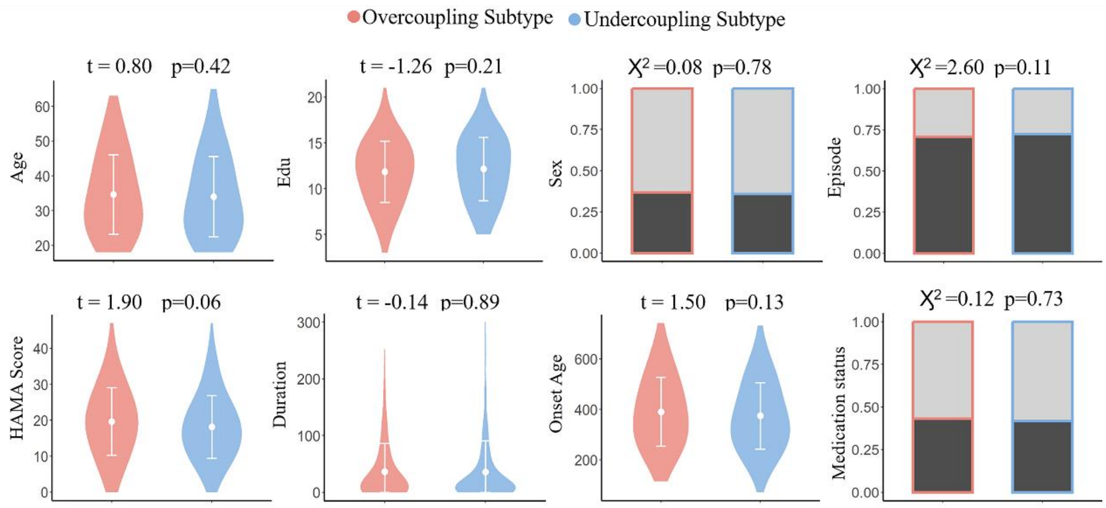


**Figure S2. Demographic and clinical differences between the two MDD subtypes.** Each point represents the mean value, and the corresponding error bar indicates the standard deviation.

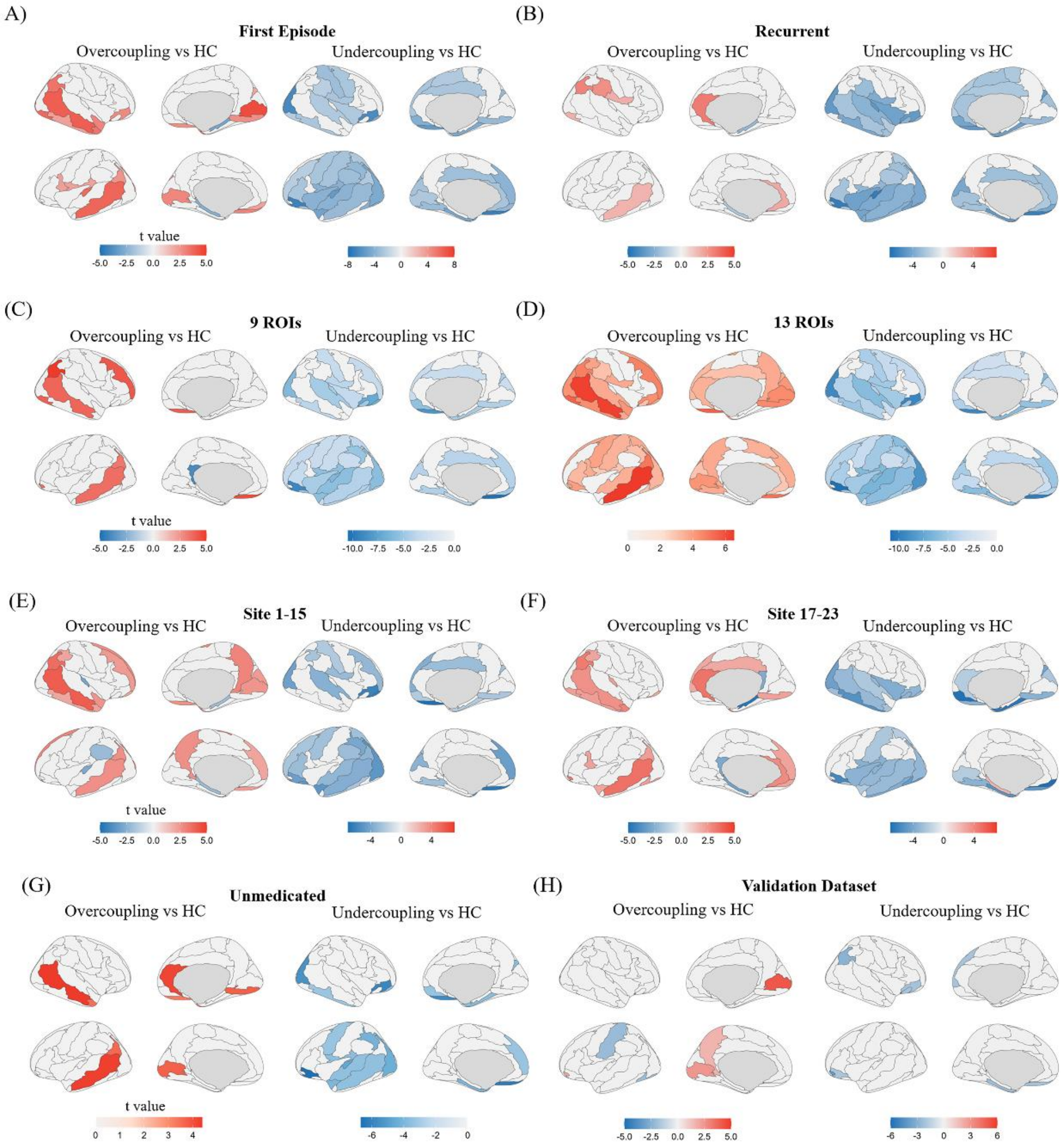


**Figure S3. Validations of MFC alteration patterns in two MDD subtypes.** Validations in first-episode and recurrent MDD patients: (A) and (B) show regional MFC differences between MDD subtype and HC in the first-episode MDD subset and in the recurrent MDD subset, respectively. Only regions that survived $p < 0.05$ are shown. Validation across different feature selection thresholds: (C) and (D) show regional MFC differences between MDD subtype and HC using 9 selected ROIs and using 13 selected ROIs. Only regions that survived FDR correction ($p < 0.05$) are shown. Half validation through splitting the REST-meta-MDD dataset: (E) and (F) show regional MFC differences between MDD subtypes and HC in two independent subsets: sites 1–15 and sites 17–23. (G) shows regional MFC differences between MDD subtype and HC in the unmedicated MDD subset. (H) shows regional MFC differences between MDD subtype and HC identified in the validation dataset. Only regions that survived $p < 0.05$ are shown.

**Table S1. Demographic and Clinical Characteristics of the Discovery and Validation datasets**

| | Discovery | | | Validation | | |
|---|---|---|---|---|---|---|
| | MDD (n = 828) | HC(n = 776) | | MDD(n = 256) | HC(n = 86) | |
| Age,Years | 34.33 ± 11.49 | 34.44 ± 13.05 | t = -0.18, p = 0.86 | 27.43 ± 6.67 | 27.98 ± 6.93 | t = -0.64, p = 0.52 |
| Gender (M/F) | 301/527 | 318/458 | $\chi^2 = 3.62$, p = 0.06 | 59/177 | 32/54 | $\chi^2 = 4.63$, p = 0.03 |
| Education,Years | 12.02 ± 3.39 | 13.61 ± 3.41 | t = -9.40, p < 0.01 | 14.41 ± 2.61 | 16.01 ± 2.64 | t = -4.87, p < 0.001 |
| Head Motion, mm | 0.07 ± 0.04 | 0.09 ± 0.03 | t = -0.30, p = 0.77 | 0.1 ± 0.02 | 0.07 ± 0.05 | t = -0.45, p = 0.68 |
| Episode | | | | | | |
| First episode | 400(48.31%) | - | - | - | - | - |
| Recurrent episode | 208(25.12%) | - | - | - | - | - |
| no recording | 220(26.57%) | - | - | - | - | - |
| Medication | | | | | | |
| drug use | 219(25.45%) | - | - | - | - | - |
| drug naïve | 298(35.99%) | - | - | - | - | - |
| no recoding | 311(37.56%) | - | - | - | - | - |
| Duration, months | 39.11 ± 61.05 (n = 684) | - | - | 10.60 ± 17.64 | - | - |
| Age at illness Onset,Years | 31.26 ± 12.38 (n = 684) | - | - | 26.71 ± 7.07 | - | - |
| HAMD | 21.18 ± 6.55 | - | - | 31.14 ± 6.50 | - | - |

| | | | | | | |
|---|---|---|---|---|---|---|
| | (n = 728) | | | | | |
| HAMA | 18.92 ± 9.11 (n = 526) | - | - | 22.29 ± 6.38 | - | - |

Abbreviation: MDD, major depressive disorder; HC, healthy controls; M, male; F, female; HAMD, Hamilton Depression Rating Scale; HAMA, Hamilton Anxiety Rating Scale.

**Table S2. Demographic and Clinical Characteristics of patients in longitudinal dataset**

| | **Baseline** | **Follow up (8 weeks)** | |
|---|---|---|---|
| Age, Years | 27.61 ± 7.80 | - | - |
| Gender (M/F) | 7/26 | - | - |
| Education, Years | 13.62 ± 3.11 | - | - |
| Duration, months | 10.36 ± 24.03 | - | - |
| Age at illness Onset, Years | 26.91 ± 7.48 | - | - |
| Motion, mm | 0.12 ± 0.04 | 0.10 ± 0.05 | $t = -0.40$, $p = 0.59$ |
| HAMD | 29.45 ± 7.67 | 14.21 ± 7.78 | $t = 10.64$, $p < 0.001$ |
| HAMA | 20.97 ± 7.539 | 9.94 ± 6.19 | $t = 10.49$, $p < 0.001$ |
| BSI-19 | 8.82 ± 7.80 | 4.64 ± 6.62 | $t = 4.18$, $p < 0.001$ |
| SHAPS | 33.21 ± 9.03 | 29.76 ± 8.30 | $t = 2.90$, $p = 0.007$ |

Abbreviation: M, male; F, female; HAMD, Hamilton Depression Rating Scale; HAMA, Hamilton Anxiety Rating Scale; BSI, Beck Scale for Suicide Ideation; SHAPS, Snaith-Hamilton Pleasure Scale.

**Table S3. ROI-based case-control differences between the two subtypes in the Discovery dataset**

| | Overcoupling vs HC | | Undercoupling vs HC | |
|---|---|---|---|---|
| ROI | t value | P(FDR) | t value | P(FDR) |
| Precentral_L | 2.14 | 0.03 | -3.24 | $1.22\times10^{-3}$ |
| Precentral_R | 0.85 | 0.39 | -2.86 | $4.27\times10^{-3}$ |
| Frontal_Sup_L | 2.27 | 0.02 | -2.17 | 0.03 |
| Frontal_Sup_R | 2.91 | $3.63\times10^{-3}$ | -2.37 | 0.02 |
| Frontal_Sup_Orb_L | 4.55 | $5.88\times10^{-6}$ | -10.28 | $<10^{-16}$ |
| Frontal_Sup_Orb_R | 2.90 | $3.75\times10^{-3}$ | -7.50 | $1.21\times10^{-13}$ |
| Frontal_Mid_L | 1.98 | 0.05 | -3.58 | $3.52\times10^{-4}$ |
| Frontal_Mid_R | 2.69 | 0.01 | -2.21 | 0.03 |
| Frontal_Mid_Orb_L | 1.77 | 0.08 | -4.80 | $1.82\times10^{-6}$ |
| Frontal_Mid_Orb_R | 2.84 | $4.64\times10^{-3}$ | -6.12 | $1.23\times10^{-9}$ |
| Frontal_Inf_Oper_L | 2.61 | 0.01 | -0.30 | 0.77 |
| Frontal_Inf_Oper_R | 0.36 | 0.72 | -0.59 | 0.55 |
| Frontal_Inf_Tri_L | 1.92 | 0.06 | -3.71 | $2.20\times10^{-4}$ |
| Frontal_Inf_Tri_R | 0.22 | 0.83 | -2.07 | 0.04 |
| Frontal_Inf_Orb_L | 2.94 | $3.38\times10^{-3}$ | -9.96 | $<10^{-16}$ |
| Frontal_Inf_Orb_R | 2.46 | 0.01 | -7.98 | $3.55\times10^{-15}$ |
| Rolandic_Oper_L | 2.13 | 0.03 | -5.73 | $1.24\times10^{-8}$ |
| Rolandic_Oper_R | 2.91 | $3.63\times10^{-3}$ | -4.66 | $3.54\times10^{-6}$ |
| Supp_Motor_Area_L | 2.87 | $4.17\times10^{-3}$ | -2.55 | 0.01 |
| Supp_Motor_Area_R | 2.92 | $3.56\times10^{-3}$ | -3.48 | $5.24\times10^{-4}$ |
| Olfactory_L | 0.82 | 0.41 | -1.73 | 0.08 |
| Olfactory_R | 1.34 | 0.18 | -2.04 | 0.04 |
| Frontal_Sup_Medial_L | 3.09 | $2.06\times10^{-3}$ | -4.85 | $1.36\times10^{-6}$ |

| | | | | |
|---|---|---|---|---|
| Frontal_Sup_Medial_R | 3.19 | $1.48 \times 10^{-3}$ | -3.81 | $1.48 \times 10^{-4}$ |
| Frontal_Med_Orb_L | 3.84 | $1.29 \times 10^{-4}$ | -4.62 | $4.22 \times 10^{-6}$ |
| Frontal_Med_Orb_R | 1.67 | 0.10 | -3.17 | $1.58 \times 10^{-3}$ |
| Rectus_L | 2.63 | 0.01 | -5.29 | $1.45 \times 10^{-7}$ |
| Rectus_R | 1.96 | 0.05 | -5.64 | $2.14 \times 10^{-8}$ |
| Insula_L | 3.10 | $1.96 \times 10^{-3}$ | -5.59 | $2.78 \times 10^{-8}$ |
| Insula_R | 3.42 | 6.55×10-4 | -3.88 | $1.12 \times 10^{-4}$ |
| Cingulum_Ant_L | 4.02 | 6.20×10-5 | -2.84 | $4.54 \times 10^{-3}$ |
| Cingulum_Ant_R | 5.04 | 5.48×10-7 | -2.94 | $3.33 \times 10^{-3}$ |
| Cingulum_Mid_L | 2.22 | 0.03 | -3.90 | $1.03 \times 10^{-4}$ |
| Cingulum_Mid_R | 3.05 | $2.34 \times 10^{-3}$ | -3.19 | $1.47 \times 10^{-3}$ |
| Cingulum_Post_L | -2.12 | 0.03 | 1.30 | 0.19 |
| Cingulum_Post_R | -1.38 | 0.17 | 0.36 | 0.72 |
| Hippocampus_L | -3.02 | $2.61 \times 10^{-3}$ | 0.30 | 0.77 |
| Hippocampus_R | -3.39 | $7.15 \times 10^{-4}$ | -0.53 | 0.59 |
| ParaHippocampal_L | 2.03 | 0.04 | -6.13 | $1.20 \times 10^{-9}$ |
| ParaHippocampal_R | 2.11 | 0.03 | -6.09 | $1.52 \times 10^{-9}$ |
| Amygdala_L | 3.15 | $1.65 \times 10^{-3}$ | -2.28 | 0.02 |
| Amygdala_R | 3.30 | $9.82 \times 10^{-4}$ | -4.25 | $2.34 \times 10^{-5}$ |
| Calcarine_L | 3.96 | $7.95 \times 10^{-5}$ | -3.76 | $1.77 \times 10^{-4}$ |
| Calcarine_R | 4.40 | $1.17 \times 10^{-5}$ | -2.39 | 0.02 |
| Cuneus_L | 2.77 | 0.01 | -4.73 | $2.55 \times 10^{-6}$ |
| Cuneus_R | 3.38 | $7.36 \times 10^{-4}$ | -3.40 | $6.84 \times 10^{-4}$ |
| Lingual_L | 2.03 | 0.04 | -4.82 | $1.63 \times 10^{-6}$ |
| Lingual_R | 3.21 | $1.37 \times 10^{-3}$ | -4.78 | $1.98 \times 10^{-6}$ |
| Occipital_Sup_L | 3.45 | $5.71 \times 10^{-4}$ | -2.77 | 0.01 |
| Occipital_Sup_R | 2.39 | 0.02 | -5.37 | $9.33 \times 10^{-8}$ |
| Occipital_Mid_L | 3.43 | $6.33 \times 10^{-4}$ | -4.93 | $9.55 \times 10^{-7}$ |
| Occipital_Mid_R | 4.05 | $5.42 \times 10^{-5}$ | -4.78 | $2.01 \times 10^{-6}$ |

| | | | | |
|---|---|---|---|---|
| Occipital_Inf_L | 0.04 | 0.97 | -6.16 | $9.66\times10^{-10}$ |
| Occipital_Inf_R | 2.22 | 0.03 | -7.94 | $4.66\times10^{-15}$ |
| Fusiform_L | 2.95 | $3.28\times10^{-3}$ | -5.41 | $7.41\times10^{-8}$ |
| Fusiform_R | 4.46 | $8.88\times10^{-6}$ | -2.64 | 0.01 |
| Postcentral_L | 2.20 | 0.03 | -4.19 | $2.96\times10^{-5}$ |
| Postcentral_R | 2.01 | 0.04 | -2.84 | $4.59\times10^{-3}$ |
| Parietal_Sup_L | 1.37 | 0.17 | -4.12 | $4.00\times10^{-5}$ |
| Parietal_Sup_R | 0.56 | 0.58 | -1.23 | 0.22 |
| Parietal_Inf_L | 1.25 | 0.21 | -4.60 | $4.72\times10^{-6}$ |
| Parietal_Inf_R | 3.06 | $2.23\times10^{-3}$ | -1.94 | 0.05 |
| SupraMarginal_L | 0.73 | 0.47 | -2.89 | $3.93\times10^{-3}$ |
| SupraMarginal_R | 2.70 | 0.01 | -2.30 | 0.02 |
| Angular_L | 3.43 | $6.15\times10^{-4}$ | -2.98 | $2.92\times10^{-3}$ |
| Angular_R | 4.96 | $8.20\times10^{-7}$ | -0.94 | 0.35 |
| Precuneus_L | 3.30 | $1.01\times10^{-3}$ | -1.22 | 0.22 |
| Precuneus_R | 3.85 | $1.22\times10^{-4}$ | -0.79 | 0.43 |
| Paracentral_Lobule_L | 2.51 | 0.01 | -1.33 | 0.18 |
| Paracentral_Lobule_R | 2.30 | 0.02 | -2.33 | 0.02 |
| Caudate_L | -6.18 | $8.85\times10^{-10}$ | 1.34 | 0.18 |
| Caudate_R | -4.31 | $1.81\times10^{-5}$ | 1.73 | 0.08 |
| Putamen_L | -0.07 | 0.95 | -1.13 | 0.26 |
| Putamen_R | 0.30 | 0.76 | -0.79 | 0.43 |
| Pallidum_L | -2.40 | 0.02 | 1.89 | 0.06 |
| Pallidum_R | -2.30 | 0.02 | 1.01 | 0.31 |
| Thalamus_L | -4.23 | $2.52\times10^{-5}$ | -1.10 | 0.27 |
| Thalamus_R | -5.45 | $5.99\times10^{-8}$ | -0.87 | 0.38 |
| Heschl_L | 3.39 | $7.26\times10^{-4}$ | -8.19 | $6.66\times10^{-16}$ |
| Heschl_R | 2.20 | 0.03 | -4.55 | $5.96\times10^{-6}$ |
| Temporal_Sup_L | 2.62 | 0.01 | -5.90 | $4.66\times10^{-9}$ |

| | | | | |
|---|---|---|---|---|
| Temporal_Sup_R | 3.01 | $2.70\times10^{-3}$ | -5.74 | $1.18\times10^{-8}$ |
| Temporal_Pole_Sup_L | 1.39 | 0.16 | -6.54 | $8.85\times10^{-11}$ |
| Temporal_Pole_Sup_R | 4.88 | $1.22\times10^{-6}$ | -2.64 | 0.01 |
| Temporal_Mid_L | 5.65 | $1.98\times10^{-8}$ | -5.77 | $1.03\times10^{-8}$ |
| Temporal_Mid_R | 5.05 | $5.10\times10^{-7}$ | -3.60 | $3.32\times10^{-4}$ |
| Temporal_Pole_Mid_L | 1.49 | 0.14 | -2.41 | 0.02 |
| Temporal_Pole_Mid_R | 3.43 | $6.27\times10^{-4}$ | -0.73 | 0.47 |
| Temporal_Inf_L | 1.97 | 0.05 | -3.64 | $2.88\times10^{-4}$ |
| Temporal_Inf_R | 3.56 | $3.86\times10^{-4}$ | -4.62 | $4.26\times10^{-6}$ |
| Cerebelum_Crus1_L | 1.65 | 0.10 | -0.44 | 0.66 |
| Cerebelum_Crus1_R | 1.52 | 0.13 | -0.22 | 0.82 |
| Cerebelum_Crus2_L | 1.68 | 0.09 | 0.29 | 0.78 |
| Cerebelum_Crus2_R | -1.13 | 0.26 | 0.58 | 0.56 |
| Cerebelum_3_L | -0.80 | 0.42 | -0.34 | 0.74 |
| Cerebelum_3_R | -1.54 | 0.12 | 0.83 | 0.41 |
| Cerebelum_4_5_L | 0.64 | 0.52 | 0.87 | 0.38 |
| Cerebelum_4_5_R | 1.22 | 0.22 | -0.65 | 0.52 |
| Cerebelum_6_L | 2.48 | 0.01 | -0.50 | 0.62 |
| Cerebelum_6_R | 2.38 | 0.02 | 0.17 | 0.87 |
| Cerebelum_7b_L | 0.78 | 0.43 | 0.95 | 0.34 |
| Cerebelum_7b_R | -0.18 | 0.86 | 0.82 | 0.41 |
| Cerebelum_8_L | -0.79 | 0.43 | 0.41 | 0.68 |
| Cerebelum_8_R | -0.38 | 0.70 | 1.93 | 0.05 |
| Cerebelum_9_L | -2.34 | 0.02 | 0.16 | 0.87 |
| Cerebelum_9_R | -1.45 | 0.15 | -1.26 | 0.21 |
| Cerebelum_10_L | -0.37 | 0.71 | 0.40 | 0.69 |
| Cerebelum_10_R | -0.36 | 0.72 | -0.51 | 0.61 |
| Vermis_1_2 | -0.96 | 0.34 | -1.14 | 0.25 |
| Vermis_3 | -2.34 | 0.02 | -0.35 | 0.73 |

| | | | | |
|---|---|---|---|---|
| Vermis_4_5 | -1.41 | 0.16 | 0.67 | 0.50 |
| Vermis_6 | -0.01 | 0.99 | -0.65 | 0.51 |
| Vermis_7 | 4.59 | $5.00\times10^{-6}$ | -0.01 | 0.99 |
| Vermis_8 | 0.46 | 0.65 | -0.65 | 0.52 |
| Vermis_9 | -2.12 | 0.03 | -2.13 | 0.03 |
| Vermis_10 | -0.16 | 0.87 | -0.78 | 0.43 |

**Table S4. ROI-based case-control differences in the Discovery dataset**

| | **MDD vs HC** | | **RMDD vs HC** | | **FEDN vs HC** | |
|---|---|---|---|---|---|---|
| **ROI** | **t value** | **P(FDR)** | **t value** | **P(FDR)** | **t value** | **P(FDR)** |
| Precentral_L | -0.63 | 0.78 | 1.74 | 0.24 | -0.69 | 0.90 |
| Precentral_R | -1.20 | 0.53 | -0.23 | 0.90 | -2.55 | 0.16 |
| Frontal_Sup_L | 0.05 | 0.98 | -0.59 | 0.76 | 0.51 | 0.93 |
| Frontal_Sup_R | 0.05 | 0.98 | -1.25 | 0.44 | -0.21 | 0.95 |
| Frontal_Sup_Orb_L | -4.04 | $3.31\times10^{-3}$ | -3.37 | 0.01 | -2.60 | 0.16 |
| Frontal_Sup_Orb_R | -3.18 | 0.02 | -3.34 | 0.01 | -1.17 | 0.72 |
| Frontal_Mid_L | -1.25 | 0.53 | -0.21 | 0.90 | -0.98 | 0.79 |
| Frontal_Mid_R | 0.33 | 0.88 | 0.36 | 0.83 | -0.14 | 0.96 |
| Frontal_Mid_Orb_L | -1.94 | 0.24 | -1.82 | 0.21 | -0.28 | 0.95 |
| Frontal_Mid_Orb_R | -2.15 | 0.18 | -2.60 | 0.06 | -0.27 | 0.95 |
| Frontal_Inf_Oper_L | 1.31 | 0.51 | 0.69 | 0.74 | 1.93 | 0.33 |
| Frontal_Inf_Oper_R | 0.11 | 0.98 | -0.37 | 0.83 | -0.22 | 0.95 |
| Frontal_Inf_Tri_L | -0.87 | 0.66 | -0.12 | 0.95 | -0.02 | 0.99 |
| Frontal_Inf_Tri_R | -0.97 | 0.62 | -0.37 | 0.83 | -0.20 | 0.95 |
| Frontal_Inf_Orb_L | -4.17 | $3.31\times10^{-3}$ | -3.88 | $4.55\times10^{-3}$ | -2.24 | 0.24 |
| Frontal_Inf_Orb_R | -3.12 | 0.02 | -3.59 | $8.41\times10^{-3}$ | -1.15 | 0.72 |

| | | | | | | |
|---|---|---|---|---|---|---|
| Rolandic_Oper_L | -2.49 | 0.10 | -1.45 | 0.35 | -0.22 | 0.95 |
| Rolandic_Oper_R | -1.22 | 0.53 | -1.00 | 0.58 | -0.10 | 0.97 |
| Supp_Motor_Area_L | 0.53 | 0.83 | -0.13 | 0.95 | 0.19 | 0.95 |
| Supp_Motor_Area_R | 0.09 | 0.98 | -0.66 | 0.74 | -0.63 | 0.90 |
| Olfactory_L | -0.74 | 0.73 | -1.42 | 0.35 | 1.35 | 0.71 |
| Olfactory_R | -0.45 | 0.83 | -0.61 | 0.76 | -0.30 | 0.95 |
| Frontal_Sup_Medial_L | -0.76 | 0.72 | -1.18 | 0.47 | -0.64 | 0.90 |
| Frontal_Sup_Medial_R | -0.45 | 0.83 | -1.19 | 0.47 | 0.08 | 0.97 |
| Frontal_Med_Orb_L | -0.43 | 0.83 | -2.57 | 0.06 | -0.17 | 0.95 |
| Frontal_Med_Orb_R | -0.69 | 0.74 | -2.35 | 0.08 | 1.28 | 0.71 |
| Rectus_L | -1.64 | 0.33 | -2.76 | 0.04 | -0.60 | 0.90 |
| Rectus_R | -2.14 | 0.18 | -3.40 | 0.01 | -0.21 | 0.95 |
| Insula_L | -1.68 | 0.33 | -2.00 | 0.16 | -0.83 | 0.84 |
| Insula_R | -1.68 | 0.33 | -2.55 | 0.06 | -0.77 | 0.88 |
| Cingulum_Ant_L | -0.06 | 0.98 | -1.84 | 0.21 | 0.67 | 0.90 |
| Cingulum_Ant_R | 0.69 | 0.74 | -1.13 | 0.50 | 2.09 | 0.29 |
| Cingulum_Mid_L | -0.93 | 0.65 | -1.44 | 0.35 | -0.82 | 0.84 |
| Cingulum_Mid_R | -0.29 | 0.88 | -0.82 | 0.68 | -0.33 | 0.95 |
| Cingulum_Post_L | -0.90 | 0.66 | -0.40 | 0.83 | -0.50 | 0.93 |
| Cingulum_Post_R | -0.60 | 0.78 | -0.53 | 0.78 | -1.21 | 0.71 |
| Hippocampus_L | -1.63 | 0.33 | -0.62 | 0.76 | -1.77 | 0.43 |
| Hippocampus_R | -2.48 | 0.10 | -1.08 | 0.53 | -2.67 | 0.16 |
| ParaHippocampal_L | -2.01 | 0.22 | -4.57 | $6.76\times10^{-4}$ | -0.39 | 0.95 |
| ParaHippocampal_R | -2.28 | 0.14 | -3.66 | $8.05\times10^{-3}$ | -0.74 | 0.89 |
| Amygdala_L | 0.61 | 0.78 | -1.49 | 0.35 | -0.17 | 0.95 |
| Amygdala_R | -1.30 | 0.51 | -1.86 | 0.20 | -0.93 | 0.79 |
| Calcarine_L | -0.06 | 0.98 | -1.45 | 0.35 | 0.43 | 0.94 |

| | | | | | | |
|---|---|---|---|---|---|---|
| Calcarine_R | 0.77 | 0.72 | -1.20 | 0.47 | 0.89 | 0.81 |
| Cuneus_L | -1.34 | 0.51 | -2.21 | 0.11 | -0.66 | 0.90 |
| Cuneus_R | -0.24 | 0.92 | -0.68 | 0.74 | 0.52 | 0.93 |
| Lingual_L | -1.89 | 0.25 | -1.39 | 0.35 | -1.08 | 0.72 |
| Lingual_R | -1.20 | 0.53 | -1.63 | 0.29 | -0.27 | 0.95 |
| Occipital_Sup_L | -0.39 | 0.85 | -0.27 | 0.87 | 0.60 | 0.90 |
| Occipital_Sup_R | -1.76 | 0.31 | -2.80 | 0.04 | -0.12 | 0.97 |
| Occipital_Mid_L | -0.86 | 0.66 | -2.63 | 0.06 | 0.71 | 0.90 |
| Occipital_Mid_R | -0.51 | 0.83 | -2.06 | 0.14 | 0.61 | 0.90 |
| Occipital_Inf_L | -3.76 | $4.08\times 10^{-3}$ | -2.77 | 0.04 | -1.57 | 0.52 |
| Occipital_Inf_R | -3.80 | $4.08\times 10^{-3}$ | -2.20 | 0.11 | -2.12 | 0.29 |
| Fusiform_L | -1.75 | 0.31 | -2.43 | 0.08 | 0.05 | 0.99 |
| Fusiform_R | 0.80 | 0.71 | 0.01 | 1.00 | 1.22 | 0.71 |
| Postcentral_L | -1.43 | 0.44 | -0.63 | 0.76 | 0.19 | 0.95 |
| Postcentral_R | -1.58 | 0.36 | -0.66 | 0.74 | -0.24 | 0.95 |
| Parietal_Sup_L | -1.95 | 0.24 | -0.76 | 0.72 | -0.74 | 0.89 |
| Parietal_Sup_R | -0.56 | 0.81 | -0.79 | 0.70 | 0.52 | 0.93 |
| Parietal_Inf_L | -2.50 | 0.10 | -0.60 | 0.76 | -1.33 | 0.71 |
| Parietal_Inf_R | 0.72 | 0.74 | 1.44 | 0.35 | 0.17 | 0.95 |
| SupraMarginal_L | -1.53 | 0.39 | -0.10 | 0.95 | -1.25 | 0.71 |
| SupraMarginal_R | -0.05 | 0.98 | 1.54 | 0.34 | -1.30 | 0.71 |
| Angular_L | -0.01 | 1.00 | -0.09 | 0.95 | 0.61 | 0.90 |
| Angular_R | 2.43 | 0.11 | 0.15 | 0.94 | 2.72 | 0.16 |
| Precuneus_L | 1.16 | 0.55 | -0.55 | 0.78 | 0.93 | 0.79 |
| Precuneus_R | 0.98 | 0.62 | -0.31 | 0.86 | 0.02 | 0.99 |
| Paracentral_Lobule_L | 0.42 | 0.83 | 0.53 | 0.78 | 0.11 | 0.97 |
| Paracentral_Lobule_R | 0.11 | 0.98 | -0.46 | 0.80 | -1.58 | 0.52 |

| | | | | | | |
|---|---|---|---|---|---|---|
| Caudate_L | -3.00 | 0.03 | -3.13 | 0.02 | -1.67 | 0.46 |
| Caudate_R | -1.65 | 0.33 | -1.42 | 0.35 | -0.91 | 0.79 |
| Putamen_L | -0.97 | 0.62 | -1.52 | 0.35 | 0.10 | 0.97 |
| Putamen_R | -0.46 | 0.83 | -1.47 | 0.35 | 0.02 | 0.99 |
| Pallidum_L | -0.45 | 0.83 | -0.94 | 0.62 | -1.98 | 0.31 |
| Pallidum_R | -0.88 | 0.66 | -0.69 | 0.74 | -2.27 | 0.24 |
| Thalamus_L | -3.25 | 0.02 | -3.42 | 0.01 | -2.79 | 0.16 |
| Thalamus_R | -3.90 | $3.81\times10^{-3}$ | -3.15 | 0.02 | -3.42 | 0.08 |
| Heschl_L | -3.16 | 0.02 | -4.00 | $4.07\times10^{-3}$ | -1.07 | 0.72 |
| Heschl_R | -1.98 | 0.23 | -2.66 | 0.05 | -0.31 | 0.95 |
| Temporal_Sup_L | -2.34 | 0.12 | -2.56 | 0.06 | -1.71 | 0.44 |
| Temporal_Sup_R | -2.05 | 0.21 | -1.69 | 0.26 | -1.15 | 0.72 |
| Temporal_Pole_Sup_L | -2.39 | 0.12 | -1.87 | 0.20 | -0.93 | 0.79 |
| Temporal_Pole_Sup_R | 0.97 | 0.62 | -0.27 | 0.87 | 0.92 | 0.79 |
| Temporal_Mid_L | -0.23 | 0.92 | -2.22 | 0.11 | 0.44 | 0.94 |
| Temporal_Mid_R | 0.97 | 0.62 | -2.03 | 0.15 | 1.72 | 0.44 |
| Temporal_Pole_Mid_L | -1.31 | 0.51 | -2.89 | 0.04 | -1.01 | 0.78 |
| Temporal_Pole_Mid_R | 0.77 | 0.72 | -2.42 | 0.08 | 2.02 | 0.30 |
| Temporal_Inf_L | -1.11 | 0.58 | -2.83 | 0.04 | -0.28 | 0.95 |
| Temporal_Inf_R | -0.59 | 0.78 | -2.34 | 0.08 | -0.24 | 0.95 |
| Cerebelum_Crus1_L | 0.50 | 0.83 | -0.54 | 0.78 | 0.59 | 0.90 |
| Cerebelum_Crus1_R | 0.33 | 0.88 | -0.87 | 0.65 | 1.89 | 0.34 |
| Cerebelum_Crus2_L | 1.01 | 0.62 | 0.18 | 0.92 | 1.23 | 0.71 |
| Cerebelum_Crus2_R | -0.31 | 0.88 | -2.24 | 0.10 | 0.66 | 0.90 |
| Cerebelum_3_L | -0.60 | 0.78 | 0.39 | 0.83 | -2.22 | 0.24 |
| Cerebelum_3_R | -0.34 | 0.88 | 0.47 | 0.80 | -1.07 | 0.72 |
| Cerebelum_4_5_L | 1.14 | 0.56 | 0.71 | 0.74 | 2.72 | 0.16 |

| | | | | | | |
|---|---|---|---|---|---|---|
| Cerebelum_4_5_R | 0.21 | 0.93 | $<10^{-16}$ | 1.00 | 1.09 | 0.72 |
| Cerebelum_6_L | 1.25 | 0.53 | 0.40 | 0.83 | 1.34 | 0.71 |
| Cerebelum_6_R | 1.48 | 0.41 | -0.43 | 0.83 | 2.01 | 0.30 |
| Cerebelum_7b_L | 1.15 | 0.55 | 0.49 | 0.79 | 0.34 | 0.95 |
| Cerebelum_7b_R | 0.44 | 0.83 | -0.90 | 0.65 | 0.21 | 0.95 |
| Cerebelum_8_L | -0.12 | 0.98 | 1.09 | 0.53 | 0.44 | 0.94 |
| Cerebelum_8_R | 1.07 | 0.60 | 0.22 | 0.90 | 0.38 | 0.95 |
| Cerebelum_9_L | -1.05 | 0.61 | -0.75 | 0.72 | -1.13 | 0.72 |
| Cerebelum_9_R | -1.65 | 0.33 | -0.87 | 0.65 | -1.21 | 0.71 |
| Cerebelum_10_L | 0.09 | 0.98 | 0.83 | 0.68 | -0.48 | 0.94 |
| Cerebelum_10_R | -0.45 | 0.83 | 0.31 | 0.86 | 0.50 | 0.93 |
| Vermis_1_2 | -1.20 | 0.53 | -1.39 | 0.35 | -2.23 | 0.24 |
| Vermis_3 | -1.87 | 0.26 | -1.31 | 0.40 | -1.08 | 0.72 |
| Vermis_4_5 | -0.70 | 0.74 | -0.71 | 0.74 | 0.85 | 0.84 |
| Vermis_6 | -0.31 | 0.88 | -2.36 | 0.08 | -0.44 | 0.94 |
| Vermis_7 | 2.94 | 0.04 | 0.04 | 0.99 | 2.60 | 0.16 |
| Vermis_8 | 0.00 | 1.00 | -0.99 | 0.58 | -0.23 | 0.95 |
| Vermis_9 | -2.61 | 0.09 | -1.44 | 0.35 | -2.43 | 0.20 |
| Vermis_10 | -0.92 | 0.65 | 0.51 | 0.79 | -1.34 | 0.71 |

**Table S5. Correlations between MFC case-control differences and meta-analytic maps for cognitive terms in Neurosynth**

| **Overcoupling Subtype** | | **Undercoupling Subtype** | |
|---|---|---|---|
| **Cognitive Term** | **r** | **Cognitive Term** | **r** |
| visual | 0.165 | music | -0.134 |
| theory mind | 0.160 | speach perception | -0.139 |
| sentences | 0.152 | audiovisual | -0.141 |
| comprehension | 0.147 | speaker | -0.147 |
| mentalizing | 0.144 | words | -0.150 |

| | | | |
|---|---|---|---|
| junction | 0.144 | conprehension | -0.151 |
| mind | 0.142 | pitch | -0.154 |
| social | 0.142 | linguistic | -0.154 |
| listening | 0.142 | primmary auditory | -0.157 |
| speech | 0.141 | spoken | -0.157 |
| motion | 0.140 | sentences | -0.159 |
| learning | -0.142 | listened | -0.165 |
| loop | -0.142 | senmantic | -0.166 |
| reward anticipation | -0.150 | language | -0.170 |
| monetary incentive | -0.161 | acoustic | -0.176 |
| incentive delay | -0.170 | visual | -0.179 |
| monetary | -0.172 | speech | -0.181 |
| reward | -0.184 | sounds | -0.185 |
| incentive | -0.187 | listening | -0.191 |
| anticipation | -0.190 | auditory | -0.203 |

**Table S6. Comparison of clinical profiles between the two subtypes**

| **HAMD17** | **t value** | **P(FDR)** |
|---|---|---|
| Depressed Mood | 0.71 | 0.88 |
| Guilt | -1.51 | 0.37 |
| Suicide | 0.27 | 0.88 |
| Early Insomia | 0.46 | 0.88 |
| Middle Insomia | 0.67 | 0.88 |
| Late Insomia | -0.86 | 0.88 |
| Work and Interest | -0.38 | 0.88 |
| Retardation | 2.93 | 0.03 |
| Agitation | 3.17 | 0.03 |
| Psychic Anxiety | 2.06 | 0.17 |

| | | |
|---|---|---|
| Somatic Anxiety | 0.58 | 0.88 |
| Gastrointetinal Symptom | 0.31 | 0.88 |
| General Somatic Symptoms | 0.45 | 0.88 |
| Genital Symptoms | -1.54 | 0.37 |
| Hypochondria | 2.71 | 0.04 |
| Weight Loss | 0.22 | 0.88 |
| Insight | 0.06 | 0.95 |

**Table S7. Nodal characteristics of symptoms in the two subtypes.**

| | **Nodal Strength** | |
|---|---|---|
| **HAMD17** | **Overcoupling** | **Undercoupling** |
| Depressed mood | 4.12 | 3.80 |
| Guilt | 1.30 | 2.19 |
| Suicide | 1.73 | 1.43 |
| Early Insomnia | 0.88 | 0.85 |
| Middle Insomnia | 3.45 | 1.12 |
| Late Insomia | 1.83 | 1.54 |
| Work and interest | 2.92 | 2.96 |
| Retaration | 1.62 | 2.13 |
| Agitation | 2.45 | 1.42 |
| Psychic Anxiety | 3.44 | 3.36 |
| Somatic Anxiety | 1.23 | 1.99 |
| Gastrointestin | 1.94 | 1.19 |
| General somatic | 2.77 | 2.82 |
| Genital | 0.21 | 0.71 |
| Hypochondria | 1.81 | 0.77 |
| Weight loss | 1.34 | 1.19 |
| Insight | 0.00 | 0.00 |

**Table S8. Enrichment analysis of PLS1 genes in GO and KEGG terms for Overcoupling Subtype**

| GO/KEGG term ID | Description | Enriched Genes (number) | log10(P-FDR) |
|---|---|---|---|
| GO:0098793 | presynapse | 75 | -19.73 |
| GO:0050804 | modulation of chemical synaptic transmission | 63 | -15.56 |
| GO:0030424 | axon | 72 | -15.05 |
| GO:0031175 | neuron projection development | 69 | -12.36 |
| GO:0120035 | regulation of plasma membrane bounded cell projection organization | 66 | -12.36 |
| GO:0016773 | phosphotransferase activity, alcohol group as acceptor | 62 | -9.64 |
| GO:0010008 | endosome membrane | 56 | -9.43 |
| GO:0048471 | perinuclear region of cytoplasm | 64 | -8.88 |
| GO:1903530 | regulation of secretion by cell | 53 | -8.68 |
| GO:0007610 | behavior | 56 | -8.41 |
| GO:0060627 | regulation of vesicle-mediated transport | 50 | -8.26 |
| GO:0019900 | kinase binding | 64 | -7.78 |
| GO:0051640 | organelle localization | 50 | -7.7 |
| GO:0022843 | voltage-gated monoatomic cation channel activity | 24 | -7.68 |
| GO:0030135 | coated vesicle | 35 | -6.83 |
| GO:0050803 | regulation of synapse structure or activity | 31 | -6.79 |
| GO:0003712 | transcription coregulator activity | 46 | -6.76 |
| GO:0031329 | regulation of cellular catabolic process | 43 | -6.61 |
| GO:0033365 | protein localization to organelle | 59 | -6.43 |

| | | | |
|---|---|---|---|
| GO:0051668 | localization within membrane | 48 | -6.42 |

**Table S9. Enrichment analysis of PLS1 genes in GO and KEGG terms for Undercoupling Subtype**

| GO/KEGG term ID | Description | Enriched Genes (number) | log10(P-FDR) |
|---|---|---|---|
| hsa05168 | Herpes simplex virus 1 infection | 56 | -9.93 |
| GO:0000902 | cell morphogenesis | 54 | -4.56 |
| GO:0030425 | dendrite | 48 | -3.84 |
| GO:0051402 | neuron apoptotic process | 16 | -3.18 |
| GO:0031252 | cell leading edge | 35 | -3.02 |
| GO:0007420 | brain development | 50 | -3.01 |
| GO:0030424 | axon | 46 | -2.96 |
| GO:0005813 | centrosome | 50 | -2.96 |
| GO:0015629 | actin cytoskeleton | 39 | -2.86 |
| GO:0072001 | renal system development | 28 | -2.82 |
| GO:0008283 | cell population proliferation | 47 | -2.44 |
| GO:0010975 | regulation of neuron projection development | 33 | -2.18 |
| GO:0003682 | chromatin binding | 42 | -2.17 |
| GO:0097190 | apoptotic signaling pathway | 26 | -2.08 |
| GO:0051301 | cell division | 36 | -1.98 |
| GO:1901615 | organic hydroxy compound metabolic process | 34 | -1.92 |
| GO:1901875 | positive regulation of post-translational protein modification | 15 | -1.92 |
| GO:0032970 | regulation of actin filament-based process | 29 | -1.85 |

| | | | |
|---|---|---|---|
| GO:0043408 | regulation of MAPK cascade | 42 | -1.83 |
| GO:1901137 | carbohydrate derivative biosynthetic process | 39 | -1.83 |

**Table S10. The correlations between MFC case-control differences and neurotransmitter densities**

| | Overcoupling | | Undercoupling | |
|---|---|---|---|---|
| **Neurotransmitter** | **r** | **Pexact-FDR** | **r** | **Pexact-FDR** |
| 5HT1a | 0.63 | $2.50\times10^{-3}$ | -0.85 | $2.12\times10^{-3}$ |
| 5HT1b | 0.43 | 0.01 | -0.55 | $2.12\times10^{-3}$ |
| 5HT2a | 0.78 | $2.50\times10^{-3}$ | -0.91 | $2.12\times10^{-3}$ |
| 5HT4 | -0.21 | $4.90\times10^{-2}$ | 0.05 | 0.58 |
| CB1 | 0.51 | $2.50\times10^{-3}$ | -0.53 | $2.12\times10^{-3}$ |
| CBF | 0.65 | $2.50\times10^{-3}$ | -0.63 | $2.12\times10^{-3}$ |
| D1 | -0.10 | 0.54 | -0.06 | 0.67 |
| D2 | -0.36 | $3.79\times10^{-3}$ | 0.19 | 0.05 |
| DAT | -0.40 | $2.12\times10^{-3}$ | 0.17 | 0.09 |
| FDOPA | -0.16 | 0.11 | 0.07 | 0.52 |
| GABAa | 0.63 | $2.50\times10^{-3}$ | -0.72 | $2.12\times10^{-3}$ |
| Kappa | 0.40 | $8.99\times10^{-3}$ | -0.51 | $2.12\times10^{-3}$ |
| MU | -0.15 | 0.43 | -0.07 | 0.67 |
| NaT | 0.01 | 0.95 | -0.18 | 0.09 |
| NMDA | -0.13 | 0.20 | -0.14 | 0.19 |
| SERT | -0.38 | $2.12\times10^{-3}$ | 0.11 | 0.28 |
| VAChT | -0.52 | $2.12\times10^{-3}$ | 0.30 | $3.79\times10^{-3}$ |
| mGluR5 | 0.51 | $2.50\times10^{-3}$ | -0.68 | $2.12\times10^{-3}$ |

**Table S11. The correlations between MFC case-control differences and cell types**

| | Overcoupling | Undercoupling |
|---|---|---|

| Cell type | r | $P_{(FDR)}$ | r | $P_{(FDR)}$ |
|---|---|---|---|---|
| Astrocytes | 0.4431 | $<10^{-3}$ | -0.2589 | 0.042 |
| Endothelial | 0.2497 | 0.020 | -0.1494 | 0.216 |
| Microglia | 0.4917 | $<10^{-3}$ | -0.3320 | 0.056 |
| Neurons | 0.3106 | 0.003 | -0.0131 | 0.529 |
| Oligodendrocytes | 0.3547 | 0.003 | -0.2694 | 0.042 |
| Oligodendrocyte precursor | 0.1067 | 0.350 | 0.1459 | 0.735 |

**Table S12. MFC differences among TRD, nTRD, and HC**

| ROI | F value | $P_{(FDR)}$ |
|---|---|---|
| Precentral_L | 2.32 | 0.16 |
| Precentral_R | 0.66 | 0.56 |
| Frontal_Sup_R | 3.15 | 0.10 |
| Frontal_Sup_Orb_L | 5.41 | 0.02 |
| Frontal_Sup_Orb_R | 2.76 | 0.12 |
| Frontal_Mid_L | 2.77 | 0.12 |
| Frontal_Mid_Orb_L | 2.11 | 0.18 |
| Frontal_Mid_Orb_R | 2.35 | 0.16 |
| Frontal_Inf_Tri_L | 5.52 | 0.02 |
| Frontal_Inf_Orb_L | 6.27 | 0.01 |
| Frontal_Inf_Orb_R | 4.07 | 0.06 |
| Rolandic_Oper_L | 2.30 | 0.16 |
| Rolandic_Oper_R | 1.81 | 0.21 |
| Supp_Motor_Area_R | 2.01 | 0.19 |
| Frontal_Sup_Medial_L | 3.24 | 0.09 |
| Frontal_Sup_Medial_R | 2.79 | 0.12 |
| Frontal_Med_Orb_L | 6.25 | 0.01 |
| Frontal_Med_Orb_R | 4.24 | 0.05 |
| Rectus_L | 2.10 | 0.18 |

| | | |
|---|---|---|
| Rectus_R | 2.18 | 0.17 |
| Insula_L | 2.09 | 0.18 |
| Insula_R | 1.73 | 0.23 |
| Cingulum_Ant_L | 1.36 | 0.30 |
| Cingulum_Ant_R | 0.52 | 0.63 |
| Cingulum_Mid_L | 1.70 | 0.23 |
| Cingulum_Mid_R | 0.48 | 0.64 |
| Hippocampus_L | 0.65 | 0.56 |
| Hippocampus_R | 1.09 | 0.38 |
| ParaHippocampal_L | 7.56 | $8.92\times10^{-3}$ |
| ParaHippocampal_R | 5.05 | 0.03 |
| Amygdala_L | 9.27 | $4.54\times10^{-3}$ |
| Amygdala_R | 6.76 | 0.01 |
| Calcarine_L | 7.37 | $8.92\times10^{-3}$ |
| Calcarine_R | 1.56 | 0.26 |
| Cuneus_L | 1.41 | 0.29 |
| Cuneus_R | 2.52 | 0.14 |
| Lingual_L | 7.47 | $8.92\times10^{-3}$ |
| Lingual_R | 6.68 | 0.01 |
| Occipital_Sup_L | 5.49 | 0.02 |
| Occipital_Sup_R | 7.13 | $9.64\times10^{-3}$ |
| Occipital_Mid_L | 8.81 | $4.54\times10^{-3}$ |
| Occipital_Mid_R | 3.84 | 0.06 |
| Occipital_Inf_L | 3.88 | 0.06 |
| Occipital_Inf_R | 3.00 | 0.11 |
| Fusiform_L | 3.89 | 0.06 |
| Fusiform_R | 3.29 | 0.09 |
| Postcentral_L | 1.25 | 0.34 |
| Postcentral_R | 1.80 | 0.21 |

| | | |
|---|---|---|
| Parietal_Sup_L | 3.95 | 0.06 |
| Parietal_Inf_L | 9.14 | $4.54\times10^{-3}$ |
| Parietal_Inf_R | 3.87 | 0.06 |
| SupraMarginal_L | 4.43 | 0.05 |
| Angular_L | 2.30 | 0.16 |
| Angular_R | 4.05 | 0.06 |
| Precuneus_L | 2.66 | 0.13 |
| Precuneus_R | 1.85 | 0.21 |
| Caudate_L | 1.15 | 0.36 |
| Caudate_R | 0.84 | 0.48 |
| Thalamus_L | 3.00 | 0.11 |
| Thalamus_R | 3.75 | 0.06 |
| Heschl_L | 0.22 | 0.81 |
| Heschl_R | 0.30 | 0.76 |
| Temporal_Sup_L | 5.24 | 0.02 |
| Temporal_Sup_R | 5.28 | 0.02 |
| Temporal_Pole_Sup_L | 2.82 | 0.12 |
| Temporal_Pole_Sup_R | 2.80 | 0.12 |
| Temporal_Mid_L | 2.30 | 0.16 |
| Temporal_Mid_R | 4.04 | 0.06 |
| Temporal_Pole_Mid_R | 1.86 | 0.21 |
| Temporal_Inf_L | 6.33 | 0.01 |
| Temporal_Inf_R | 3.49 | 0.08 |
| Vermis_7 | 0.08 | 0.92 |

**Table S13. MFC differences between pairwise groups**

| | TRD vs nTRD | TRD vs HC | nTRD vs HC |
|---|---|---|---|

| ROI | t value | P(FDR) | t value | P(FDR) | t value | P(FDR) |
|---|---|---|---|---|---|---|
| Frontal_Sup_Orb_L | -2.27 | 0.03 | -3.33 | $3.46\times10^{-3}$ | -1.50 | 0.35 |
| Frontal_Inf_Tri_L | -3.02 | $3.86\times10^{-3}$ | -0.87 | 0.39 | 2.23 | 0.19 |
| Frontal_Inf_Orb_L | -2.95 | $4.16\times10^{-3}$ | -3.49 | $2.96\times10^{-3}$ | -0.67 | 0.80 |
| Frontal_Med_Orb_R | -1.61 | 0.11 | -3.65 | $2.23\times10^{-3}$ | -2.39 | 0.19 |
| ParaHippocampal_L | -3.21 | $2.60\times10^{-3}$ | -3.67 | $2.23\times10^{-3}$ | -0.94 | 0.60 |
| ParaHippocampal_R | -3.03 | $3.86\times10^{-3}$ | -2.62 | 0.01 | 0.04 | 0.97 |
| Amygdala_L | -2.76 | $6.94\times10^{-3}$ | -4.32 | $5.33\times10^{-4}$ | -2.18 | 0.19 |
| Amygdala_R | -3.40 | $1.68\times10^{-3}$ | -3.19 | $4.64\times10^{-3}$ | -0.30 | 0.89 |
| Calcarine_L | -3.63 | $1.32\times10^{-3}$ | -3.36 | $3.46\times10^{-3}$ | -0.10 | 0.97 |
| Lingual_L | -3.69 | $1.30\times10^{-3}$ | -3.09 | $5.59\times10^{-3}$ | 0.35 | 0.89 |
| Lingual_R | -3.44 | $1.68\times10^{-3}$ | -2.96 | $7.61\times10^{-3}$ | 0.26 | 0.89 |
| Occipital_Sup_L | -3.24 | $2.57\times10^{-3}$ | -1.72 | 0.10 | 1.45 | 0.35 |
| Occipital_Sup_R | -3.70 | $1.30\times10^{-3}$ | -1.89 | 0.08 | 1.72 | 0.35 |
| Occipital_Mid_L | -4.30 | $3.13\times10^{-4}$ | -2.42 | 0.02 | 1.47 | 0.35 |
| Parietal_Inf_L | -4.23 | $3.13\times10^{-4}$ | -2.91 | $7.84\times10^{-3}$ | 1.02 | 0.59 |
| SupraMarginal_L | -3.02 | $3.86\times10^{-3}$ | -1.16 | 0.26 | 1.59 | 0.35 |
| Temporal_Sup_L | -3.43 | $1.68\times10^{-3}$ | -1.59 | 0.13 | 1.37 | 0.36 |
| Temporal_Sup_R | -2.95 | $4.16\times10^{-3}$ | -2.88 | $7.84\times10^{-3}$ | -0.41 | 0.89 |
| Temporal_Inf_L | -3.54 | $1.55\times10^{-3}$ | -2.67 | 0.01 | 0.28 | 0.89 |

**Table S14. ROI-based case-control difference in the half validation**

| | MDD vs NC(Site 1-15) | | | MDD vs NC(Site 17-23) | | |
|---|---|---|---|---|---|---|
| ROI | t value | P | P(FDR) | t value | P | P(FDR) |
| Precentral_L | -1.06 | 0.29 | 0.79 | 0.27 | 0.79 | 0.93 |
| Precentral_R | -0.45 | 0.65 | 0.84 | -1.17 | 0.24 | 0.63 |
| Frontal_Sup_L | 0.69 | 0.49 | 0.84 | -0.52 | 0.60 | 0.87 |
| Frontal_Sup_R | 0.81 | 0.42 | 0.84 | -0.58 | 0.56 | 0.86 |
| Frontal_Sup_Orb_L | -2.28 | 0.02 | 0.24 | -3.01 | $2.70\times10^{-3}$ | 0.05 |

| | | | | | | |
|---|---|---|---|---|---|---|
| Frontal_Sup_Orb_R | -2.78 | $5.52\times10^{-3}$ | 0.09 | -1.74 | 0.08 | 0.42 |
| Frontal_Mid_L | -0.74 | 0.46 | 0.84 | -0.70 | 0.48 | 0.83 |
| Frontal_Mid_R | -0.47 | 0.64 | 0.84 | 1.00 | 0.32 | 0.75 |
| Frontal_Mid_Orb_L | -1.92 | 0.06 | 0.35 | -0.89 | 0.37 | 0.78 |
| Frontal_Mid_Orb_R | -2.43 | 0.02 | 0.19 | -0.61 | 0.54 | 0.85 |
| Frontal_Inf_Oper_L | -0.14 | 0.89 | 0.94 | 1.95 | 0.05 | 0.37 |
| Frontal_Inf_Oper_R | 0.00 | 1.00 | 1.00 | 0.01 | 0.99 | 0.99 |
| Frontal_Inf_Tri_L | -1.27 | 0.21 | 0.68 | 0.06 | 0.95 | 0.98 |
| Frontal_Inf_Tri_R | -0.44 | 0.66 | 0.85 | -0.84 | 0.40 | 0.78 |
| Frontal_Inf_Orb_L | -2.25 | 0.02 | 0.24 | -3.47 | $5.52\times10^{-4}$ | 0.03 |
| Frontal_Inf_Orb_R | -2.48 | 0.01 | 0.19 | -1.86 | 0.06 | 0.39 |
| Rolandic_Oper_L | -2.05 | 0.04 | 0.32 | -1.52 | 0.13 | 0.48 |
| Rolandic_Oper_R | -1.75 | 0.08 | 0.40 | -0.08 | 0.94 | 0.98 |
| Supp_Motor_Area_L | 0.24 | 0.81 | 0.94 | 0.67 | 0.50 | 0.83 |
| Supp_Motor_Area_R | -0.04 | 0.97 | 0.98 | 0.75 | 0.46 | 0.81 |
| Olfactory_L | -0.49 | 0.62 | 0.84 | -0.67 | 0.50 | 0.83 |
| Olfactory_R | -0.54 | 0.59 | 0.84 | -0.13 | 0.89 | 0.96 |
| Frontal_Sup_Medial_L | -1.30 | 0.20 | 0.67 | 0.28 | 0.78 | 0.93 |
| Frontal_Sup_Medial_R | -1.14 | 0.26 | 0.72 | 0.59 | 0.55 | 0.85 |
| Frontal_Med_Orb_L | -0.25 | 0.80 | 0.94 | -0.48 | 0.63 | 0.87 |
| Frontal_Med_Orb_R | -0.51 | 0.61 | 0.84 | -0.33 | 0.74 | 0.92 |
| Rectus_L | -0.21 | 0.83 | 0.94 | -3.13 | $1.83\times10^{-3}$ | 0.05 |
| Rectus_R | -0.64 | 0.52 | 0.84 | -3.08 | $2.14\times10^{-3}$ | 0.05 |
| Insula_L | -0.91 | 0.36 | 0.84 | -1.52 | 0.13 | 0.48 |
| Insula_R | -0.59 | 0.56 | 0.84 | -1.73 | 0.08 | 0.42 |
| Cingulum_Ant_L | -0.53 | 0.59 | 0.84 | 0.48 | 0.63 | 0.87 |
| Cingulum_Ant_R | 0.16 | 0.88 | 0.94 | 0.87 | 0.38 | 0.78 |
| Cingulum_Mid_L | -1.96 | 0.05 | 0.35 | 0.23 | 0.82 | 0.93 |
| Cingulum_Mid_R | -1.75 | 0.08 | 0.40 | 1.32 | 0.19 | 0.53 |

| | | | | | | |
|---|---|---|---|---|---|---|
| Cingulum_Post_L | 0.73 | 0.46 | 0.84 | -1.85 | 0.06 | 0.39 |
| Cingulum_Post_R | 0.85 | 0.40 | 0.84 | -2.36 | 0.02 | 0.24 |
| Hippocampus_L | -1.74 | 0.08 | 0.40 | -0.39 | 0.69 | 0.91 |
| Hippocampus_R | -1.38 | 0.17 | 0.63 | -2.05 | 0.04 | 0.31 |
| ParaHippocampal_L | -1.31 | 0.19 | 0.67 | -2.25 | 0.02 | 0.24 |
| ParaHippocampal_R | -1.19 | 0.23 | 0.72 | -3.02 | $2.61\times10^{-3}$ | 0.05 |
| Amygdala_L | -0.14 | 0.89 | 0.94 | 1.17 | 0.24 | 0.63 |
| Amygdala_R | -1.07 | 0.28 | 0.79 | -0.89 | 0.37 | 0.78 |
| Calcarine_L | -0.18 | 0.86 | 0.94 | 0.03 | 0.98 | 0.99 |
| Calcarine_R | 0.57 | 0.57 | 0.84 | 0.26 | 0.79 | 0.93 |
| Cuneus_L | -1.87 | 0.06 | 0.36 | -0.13 | 0.89 | 0.96 |
| Cuneus_R | 0.10 | 0.92 | 0.95 | -0.62 | 0.54 | 0.85 |
| Lingual_L | -1.02 | 0.31 | 0.79 | -1.69 | 0.09 | 0.42 |
| Lingual_R | -0.93 | 0.35 | 0.84 | -0.84 | 0.40 | 0.78 |
| Occipital_Sup_L | -0.12 | 0.90 | 0.94 | -0.35 | 0.73 | 0.92 |
| Occipital_Sup_R | -1.03 | 0.30 | 0.79 | -1.42 | 0.16 | 0.50 |
| Occipital_Mid_L | 0.55 | 0.58 | 0.84 | -1.90 | 0.06 | 0.39 |
| Occipital_Mid_R | 0.03 | 0.97 | 0.98 | -0.83 | 0.41 | 0.78 |
| Occipital_Inf_L | -3.29 | $1.03\times10^{-3}$ | 0.04 | -2.08 | 0.04 | 0.31 |
| Occipital_Inf_R | -3.31 | $9.75\times10^{-4}$ | 0.04 | -2.29 | 0.02 | 0.24 |
| Fusiform_L | -1.14 | 0.26 | 0.72 | -1.54 | 0.12 | 0.48 |
| Fusiform_R | 0.66 | 0.51 | 0.84 | 0.12 | 0.90 | 0.96 |
| Postcentral_L | -0.60 | 0.55 | 0.84 | -1.41 | 0.16 | 0.50 |
| Postcentral_R | -0.75 | 0.45 | 0.84 | -0.87 | 0.38 | 0.78 |
| Parietal_Sup_L | -1.75 | 0.08 | 0.40 | -0.88 | 0.38 | 0.78 |
| Parietal_Sup_R | -0.76 | 0.45 | 0.84 | -0.05 | 0.96 | 0.98 |
| Parietal_Inf_L | -2.95 | $3.26\times10^{-3}$ | 0.06 | -0.43 | 0.67 | 0.89 |
| Parietal_Inf_R | -0.06 | 0.95 | 0.97 | 1.14 | 0.26 | 0.63 |
| SupraMarginal_L | -3.24 | $1.26\times10^{-3}$ | 0.04 | 1.36 | 0.18 | 0.52 |

| | | | | | | |
|---|---|---|---|---|---|---|
| SupraMarginal_R | -0.86 | 0.39 | 0.84 | 0.66 | 0.51 | 0.83 |
| Angular_L | -0.64 | 0.52 | 0.84 | 0.75 | 0.45 | 0.81 |
| Angular_R | 2.13 | 0.03 | 0.30 | 1.46 | 0.14 | 0.49 |
| Precuneus_L | 0.57 | 0.57 | 0.84 | 0.79 | 0.43 | 0.81 |
| Precuneus_R | 0.50 | 0.62 | 0.84 | 1.41 | 0.16 | 0.50 |
| Paracentral_Lobule_L | 0.22 | 0.82 | 0.94 | 0.52 | 0.60 | 0.87 |
| Paracentral_Lobule_R | 0.16 | 0.87 | 0.94 | -0.50 | 0.62 | 0.87 |
| Caudate_L | -1.95 | 0.05 | 0.35 | -2.24 | 0.03 | 0.24 |
| Caudate_R | -0.57 | 0.57 | 0.84 | -1.69 | 0.09 | 0.42 |
| Putamen_L | -0.73 | 0.47 | 0.84 | -0.61 | 0.54 | 0.85 |
| Putamen_R | 0.29 | 0.77 | 0.92 | -0.73 | 0.46 | 0.82 |
| Pallidum_L | 0.27 | 0.79 | 0.94 | -0.57 | 0.57 | 0.86 |
| Pallidum_R | 0.97 | 0.33 | 0.82 | -1.84 | 0.07 | 0.39 |
| Thalamus_L | -1.52 | 0.13 | 0.53 | -2.77 | $5.73\times10^{-3}$ | 0.09 |
| Thalamus_R | -1.60 | 0.11 | 0.49 | -3.62 | $3.15\times10^{-4}$ | 0.03 |
| Heschl_L | -3.21 | $1.39\times10^{-3}$ | 0.04 | -1.61 | 0.11 | 0.48 |
| Heschl_R | -2.07 | 0.04 | 0.32 | -0.55 | 0.58 | 0.87 |
| Temporal_Sup_L | -1.90 | 0.06 | 0.35 | -1.57 | 0.12 | 0.48 |
| Temporal_Sup_R | -1.72 | 0.09 | 0.40 | -1.14 | 0.25 | 0.63 |
| Temporal_Pole_Sup_L | -3.09 | $2.07\times10^{-3}$ | 0.05 | -0.05 | 0.96 | 0.98 |
| Temporal_Pole_Sup_R | 0.71 | 0.48 | 0.84 | 0.26 | 0.79 | 0.93 |
| Temporal_Mid_L | -0.75 | 0.46 | 0.84 | 0.35 | 0.73 | 0.92 |
| Temporal_Mid_R | 1.38 | 0.17 | 0.63 | -0.15 | 0.88 | 0.96 |
| Temporal_Pole_Mid_L | -0.70 | 0.48 | 0.84 | -2.16 | 0.03 | 0.28 |
| Temporal_Pole_Mid_R | 0.55 | 0.58 | 0.84 | 0.21 | 0.84 | 0.93 |
| Temporal_Inf_L | -0.40 | 0.69 | 0.87 | -1.56 | 0.12 | 0.48 |
| Temporal_Inf_R | 0.66 | 0.51 | 0.84 | -1.77 | 0.08 | 0.42 |
| Cerebelum_Crus1_L | 0.63 | 0.53 | 0.84 | 0.32 | 0.75 | 0.92 |
| Cerebelum_Crus1_R | 0.74 | 0.46 | 0.84 | -0.48 | 0.63 | 0.87 |

| | | | | | | |
|---|---|---|---|---|---|---|
| Cerebelum_Crus2_L | 0.51 | 0.61 | 0.84 | 0.90 | 0.37 | 0.78 |
| Cerebelum_Crus2_R | -0.32 | 0.75 | 0.90 | -0.25 | 0.80 | 0.93 |
| Cerebelum_3_L | -1.17 | 0.24 | 0.72 | 0.43 | 0.67 | 0.89 |
| Cerebelum_3_R | -0.57 | 0.57 | 0.84 | 0.67 | 0.50 | 0.83 |
| Cerebelum_4_5_L | 0.13 | 0.90 | 0.94 | 1.48 | 0.14 | 0.49 |
| Cerebelum_4_5_R | 0.53 | 0.60 | 0.84 | -0.10 | 0.92 | 0.97 |
| Cerebelum_6_L | 0.15 | 0.88 | 0.94 | 1.34 | 0.18 | 0.52 |
| Cerebelum_6_R | 0.48 | 0.63 | 0.84 | 1.25 | 0.21 | 0.59 |
| Cerebelum_7b_L | 0.89 | 0.37 | 0.84 | 0.76 | 0.45 | 0.81 |
| Cerebelum_7b_R | 0.45 | 0.65 | 0.84 | 0.38 | 0.70 | 0.91 |
| Cerebelum_8_L | -0.34 | 0.73 | 0.89 | 0.43 | 0.67 | 0.89 |
| Cerebelum_8_R | 1.22 | 0.22 | 0.72 | 0.20 | 0.84 | 0.93 |
| Cerebelum_9_L | -0.38 | 0.70 | 0.87 | -1.51 | 0.13 | 0.48 |
| Cerebelum_9_R | -0.99 | 0.32 | 0.81 | -1.14 | 0.25 | 0.63 |
| Cerebelum_10_L | -0.85 | 0.40 | 0.84 | 0.78 | 0.44 | 0.81 |
| Cerebelum_10_R | 0.37 | 0.71 | 0.88 | -1.05 | 0.30 | 0.71 |
| Vermis_1_2 | -1.42 | 0.15 | 0.62 | 0.28 | 0.78 | 0.93 |
| Vermis_3 | -1.58 | 0.11 | 0.49 | -0.93 | 0.35 | 0.78 |
| Vermis_4_5 | -0.53 | 0.60 | 0.84 | -0.33 | 0.74 | 0.92 |
| Vermis_6 | -1.17 | 0.24 | 0.72 | 0.95 | 0.34 | 0.78 |
| Vermis_7 | 1.30 | 0.19 | 0.67 | 2.37 | 0.02 | 0.24 |
| Vermis_8 | 0.22 | 0.83 | 0.94 | -0.21 | 0.84 | 0.93 |
| Vermis_9 | -2.41 | 0.02 | 0.19 | -1.34 | 0.18 | 0.52 |
| Vermis_10 | -0.78 | 0.44 | 0.84 | -0.21 | 0.84 | 0.93 |

**Table S15. ROI-based case-control difference in the Validation dataset**

| | MDD vs NC | Subtype1 vs NC | Subtype2 vs NC |
|---|---|---|---|

| ROI | t value | P | P(FDR) | t value | P | P(FDR) | t value | P | P(FDR) |
|---|---|---|---|---|---|---|---|---|---|
| Precentral_L | 1.81 | 0.07 | 0.39 | -1.43 | 0.15 | 0.21 | 1.44 | 0.15 | 0.57 |
| Precentral_R | 1.39 | 0.16 | 0.60 | -1.73 | 0.09 | 0.42 | 0.97 | 0.33 | 0.71 |
| Frontal_Sup_L | -0.79 | 0.43 | 0.80 | 0.62 | 0.54 | 0.84 | -0.14 | 0.89 | 0.95 |
| Frontal_Sup_R | 0.09 | 0.93 | 0.97 | -0.42 | 0.67 | 0.90 | 0.27 | 0.79 | 0.88 |
| Frontal_Sup_Orb_L | -2.35 | 0.02 | 0.22 | 1.75 | 0.08 | 0.42 | -2.20 | 0.03 | 0.28 |
| Frontal_Sup_Orb_R | -1.64 | 0.10 | 0.47 | 1.10 | 0.27 | 0.67 | -1.47 | 0.14 | 0.57 |
| Frontal_Mid_L | 0.82 | 0.41 | 0.80 | -1.44 | 0.15 | 0.51 | 0.76 | 0.45 | 0.78 |
| Frontal_Mid_R | 0.43 | 0.67 | 0.88 | -1.12 | 0.26 | 0.67 | 0.11 | 0.91 | 0.95 |
| Frontal_Mid_Orb_L | -0.44 | 0.66 | 0.88 | -0.13 | 0.90 | 0.97 | -0.94 | 0.35 | 0.71 |
| Frontal_Mid_Orb_R | -0.99 | 0.32 | 0.77 | 0.24 | 0.81 | 0.95 | -1.43 | 0.15 | 0.57 |
| Frontal_Inf_Oper_L | 1.45 | 0.15 | 0.57 | -1.73 | 0.09 | 0.42 | 1.27 | 0.21 | 0.58 |
| Frontal_Inf_Oper_R | 1.93 | 0.05 | 0.36 | -2.69 | $7.85 \times 10^{-3}$ | 0.15 | 1.05 | 0.29 | 0.66 |
| Frontal_Inf_Tri_L | 1.27 | 0.21 | 0.67 | -1.87 | 0.06 | 0.42 | 0.62 | 0.54 | 0.83 |
| Frontal_Inf_Tri_R | 0.11 | 0.91 | 0.97 | -1.03 | 0.30 | 0.68 | -0.36 | 0.72 | 0.87 |
| Frontal_Inf_Orb_L | -1.88 | 0.06 | 0.36 | 1.13 | 0.26 | 0.67 | -2.02 | 0.04 | 0.30 |
| Frontal_Inf_Orb_R | -2.16 | 0.03 | 0.31 | 1.09 | 0.28 | 0.67 | -2.17 | 0.03 | 0.28 |
| Rolandic_Oper_L | 0.35 | 0.72 | 0.89 | -1.71 | 0.09 | 0.42 | -0.50 | 0.62 | 0.87 |

| | | | | | | | | | |
|---|---|---|---|---|---|---|---|---|---|
| Rolandic_Oper_R | 1.06 | 0.29 | 0.76 | -2.05 | 0.04 | 0.40 | 0.40 | 0.69 | 0.87 |
| Supp_Motor_Area_L | 0.16 | 0.87 | 0.94 | -1.11 | 0.27 | 0.67 | -0.52 | 0.60 | 0.87 |
| Supp_Motor_Area_R | 0.41 | 0.69 | 0.88 | -1.67 | 0.10 | 0.42 | -0.59 | 0.56 | 0.84 |
| Olfactory_L | -1.65 | 0.10 | 0.47 | 0.88 | 0.38 | 0.71 | -2.20 | 0.03 | 0.28 |
| Olfactory_R | -0.44 | 0.66 | 0.88 | -0.13 | 0.90 | 0.97 | -1.01 | 0.31 | 0.68 |
| Frontal_Sup_Medial_L | -1.37 | 0.17 | 0.60 | 0.82 | 0.41 | 0.74 | -1.28 | 0.20 | 0.58 |
| Frontal_Sup_Medial_R | -2.37 | 0.02 | 0.22 | 1.63 | 0.11 | 0.42 | -2.12 | 0.04 | 0.29 |
| Frontal_Med_Orb_L | -3.70 | $2.51\times10^{-4}$ | 0.03 | 1.73 | 0.09 | 0.42 | -5.15 | $6.06\times10^{-7}$ | $7.03\times10^{-5}$ |
| Frontal_Med_Orb_R | -2.88 | $4.24\times10^{-3}$ | 0.08 | 0.64 | 0.52 | 0.84 | -4.45 | $1.42\times10^{-5}$ | $8.22\times10^{-4}$ |
| Rectus_L | -0.18 | 0.85 | 0.94 | -0.94 | 0.35 | 0.70 | -1.26 | 0.21 | 0.58 |
| Rectus_R | -0.23 | 0.82 | 0.93 | -1.43 | 0.15 | 0.51 | -1.44 | 0.15 | 0.57 |
| Insula_L | 0.64 | 0.52 | 0.88 | -1.00 | 0.05 | 0.40 | -0.66 | 0.51 | 0.80 |
| Insula_R | 0.50 | 0.62 | 0.88 | -1.81 | 0.07 | 0.42 | -0.47 | 0.64 | 0.87 |
| Cingulum_Ant_L | -0.80 | 0.43 | 0.80 | -0.46 | 0.65 | 0.90 | -1.77 | 0.08 | 0.45 |
| Cingulum_Ant_R | 0.07 | 0.95 | 0.97 | -0.92 | 0.36 | 0.71 | -0.82 | 0.41 | 0.76 |
| Cingulum_Mid_L | -0.31 | 0.75 | 0.91 | -0.11 | 0.92 | 0.97 | -0.74 | 0.46 | 0.78 |
| Cingulum_Mid_R | 0.43 | 0.67 | 0.88 | -1.16 | 0.25 | 0.67 | -0.71 | 0.48 | 0.79 |
| Cingulum_Post_ | 1.72 | 0.09 | 0.44 | -1.65 | 0.10 | 0.42 | 1.35 | 0.18 | 0.57 |

| | | | | | | | | | |
|---|---|---|---|---|---|---|---|---|---|
| L | | | | | | | | | |
| Cingulum_Post_R | 1.29 | 0.20 | 0.67 | -1.01 | 0.31 | 0.68 | 1.37 | 0.17 | 0.57 |
| Hippocampus_L | 0.56 | 0.58 | 0.88 | -0.60 | 0.55 | 0.85 | 0.42 | 0.67 | 0.87 |
| Hippocampus_R | 0.00 | 1.00 | 1.00 | 0.17 | 0.87 | 0.97 | 0.28 | 0.78 | 0.88 |
| ParaHippocampal_L | -1.89 | 0.06 | 0.36 | 0.88 | 0.38 | 0.71 | -2.40 | 0.02 | 0.21 |
| ParaHippocampal_R | -0.76 | 0.45 | 0.82 | -0.63 | 0.53 | 0.84 | -1.76 | 0.08 | 0.45 |
| Amygdala_L | -3.26 | $1.22\times10^{-3}$ | 0.04 | 1.90 | 0.06 | 0.42 | -3.99 | $9.21\times10^{-5}$ | $3.56\times10^{-3}$ |
| Amygdala_R | -1.26 | 0.21 | 0.67 | -0.05 | 0.96 | 0.97 | -2.09 | 0.04 | 0.29 |
| Calcarine_L | -2.01 | 0.05 | 0.35 | 2.49 | 0.01 | 0.20 | -1.23 | 0.22 | 0.58 |
| Calcarine_R | -2.00 | 0.05 | 0.35 | 4.20 | $4.16\times10^{-5}$ | $4.83\times10^{-3}$ | 0.37 | 0.71 | 0.87 |
| Cuneus_L | -1.02 | 0.31 | 0.77 | 1.03 | 0.30 | 0.68 | -0.77 | 0.44 | 0.78 |
| Cuneus_R | -0.39 | 0.70 | 0.88 | 0.40 | 0.69 | 0.91 | -0.33 | 0.74 | 0.87 |
| Lingual_L | -0.92 | 0.36 | 0.79 | 0.66 | 0.51 | 0.84 | -0.73 | 0.47 | 0.78 |
| Lingual_R | -1.14 | 0.25 | 0.75 | 1.05 | 0.30 | 0.68 | -0.74 | 0.46 | 0.78 |
| Occipital_Sup_L | 0.07 | 0.94 | 0.97 | -0.51 | 0.61 | 0.88 | -0.11 | 0.91 | 0.95 |
| Occipital_Sup_R | 0.06 | 0.95 | 0.97 | -0.78 | 0.44 | 0.77 | 0.00 | 1.00 | 1.00 |
| Occipital_Mid_L | -0.56 | 0.58 | 0.88 | -0.98 | 0.33 | 0.68 | -1.75 | 0.08 | 0.45 |
| Occipital_Mid_R | -2.16 | 0.03 | 0.31 | 0.66 | 0.51 | 0.84 | -2.88 | $4.43\times10^{-3}$ | 0.07 |
| Occipital_Inf_L | -0.86 | 0.39 | 0.80 | 0.14 | 0.89 | 0.97 | -1.39 | 0.17 | 0.57 |
| Occipital_Inf_R | -1.09 | 0.28 | 0.76 | 0.31 | 0.76 | 0.95 | -1.18 | 0.24 | 0.59 |

| | | | | | | | | | |
|---|---|---|---|---|---|---|---|---|---|
| Fusiform_L | 0.68 | 0.50 | 0.88 | -1.99 | 0.05 | 0.40 | -0.41 | 0.68 | 0.87 |
| Fusiform_R | -0.62 | 0.53 | 0.88 | -0.53 | 0.60 | 0.88 | -1.63 | 0.11 | 0.53 |
| Postcentral_L | 1.00 | 0.32 | 0.77 | -2.07 | 0.04 | 0.40 | 0.35 | 0.73 | 0.87 |
| Postcentral_R | 0.27 | 0.79 | 0.92 | -0.89 | 0.38 | 0.71 | -0.13 | 0.89 | 0.95 |
| Parietal_Sup_L | -0.82 | 0.41 | 0.80 | 0.58 | 0.56 | 0.86 | -0.54 | 0.59 | 0.87 |
| Parietal_Sup_R | 0.18 | 0.85 | 0.94 | -0.57 | 0.57 | 0.86 | 0.35 | 0.73 | 0.87 |
| Parietal_Inf_L | -0.20 | 0.84 | 0.94 | -0.49 | 0.62 | 0.89 | -0.68 | 0.50 | 0.80 |
| Parietal_Inf_R | 0.43 | 0.67 | 0.88 | -0.93 | 0.35 | 0.71 | -0.10 | 0.92 | 0.95 |
| SupraMarginal_L | 1.09 | 0.28 | 0.76 | -1.19 | 0.24 | 0.67 | 0.93 | 0.35 | 0.71 |
| SupraMarginal_R | 0.20 | 0.84 | 0.94 | -0.26 | 0.79 | 0.95 | 0.14 | 0.89 | 0.95 |
| Angular_L | 0.85 | 0.40 | 0.80 | -1.34 | 0.18 | 0.57 | 0.28 | 0.78 | 0.88 |
| Angular_R | -1.92 | 0.06 | 0.36 | 0.28 | 0.78 | 0.95 | -2.93 | $3.74\times10^{-3}$ | 0.07 |
| Precuneus_L | -1.19 | 0.24 | 0.74 | 1.66 | 0.10 | 0.42 | -0.61 | 0.54 | 0.83 |
| Precuneus_R | -0.33 | 0.74 | 0.90 | 0.35 | 0.73 | 0.94 | -0.35 | 0.73 | 0.87 |
| Paracentral_Lobule_L | -0.23 | 0.82 | 0.93 | -0.19 | 0.85 | 0.97 | -0.39 | 0.70 | 0.87 |
| Paracentral_Lobule_R | -1.50 | 0.13 | 0.57 | 1.49 | 0.14 | 0.51 | -0.94 | 0.35 | 0.71 |
| Caudate_L | 1.03 | 0.30 | 0.77 | -1.45 | 0.15 | 0.51 | 0.47 | 0.64 | 0.87 |
| Caudate_R | 0.03 | 0.97 | 0.98 | -0.45 | 0.65 | 0.90 | -0.43 | 0.67 | 0.87 |
| Putamen_L | -1.46 | 0.14 | 0.57 | 0.99 | 0.32 | 0.68 | -1.90 | 0.06 | 0.38 |
| Putamen_R | 0.30 | 0.77 | 0.92 | -1.55 | 0.12 | 0.48 | -1.21 | 0.23 | 0.58 |
| Pallidum_L | 2.54 | 0.01 | 0.17 | -3.52 | $5.48\times10^{-4}$ | 0.02 | 0.91 | 0.36 | 0.71 |
| Pallidum_R | 3.45 | $6.38\times10^{-4}$ | 0.03 | -3.99 | $9.27\times10^{-5}$ | $5.37\times10^{-3}$ | 2.38 | 0.02 | 0.21 |

| | | | | | | | | | |
|---|---|---|---|---|---|---|---|---|---|
| Thalamus_L | 3.42 | 7.04 ×10-4 | 0.03 | -2.71 | 7.34 ×10-3 | 0.15 | 3.58 | 4.35× 10^-4 | 0.01 |
| Thalamus_R | 2.65 | 8.44 ×10-3 | 0.14 | -1.70 | 0.09 | 0.42 | 3.09 | 2.27× 10-3 | 0.05 |
| Heschl_L | 0.58 | 0.56 | 0.88 | -1.05 | 0.30 | 0.68 | 0.11 | 0.91 | 0.95 |
| Heschl_R | -0.53 | 0.60 | 0.88 | -0.63 | 0.53 | 0.84 | -1.36 | 0.18 | 0.57 |
| Temporal_Sup_L | 0.62 | 0.54 | 0.88 | -1.84 | 0.07 | 0.42 | -0.16 | 0.87 | 0.95 |
| Temporal_Sup_R | -1.08 | 0.28 | 0.76 | -0.08 | 0.94 | 0.97 | -1.50 | 0.13 | 0.57 |
| Temporal_Pole_Sup_L | -0.40 | 0.69 | 0.88 | -0.34 | 0.74 | 0.94 | -0.75 | 0.45 | 0.78 |
| Temporal_Pole_Sup_R | -0.10 | 0.92 | 0.97 | -0.36 | 0.72 | 0.94 | -0.58 | 0.56 | 0.84 |
| Temporal_Mid_L | -0.60 | 0.55 | 0.88 | -0.29 | 0.77 | 0.95 | -1.06 | 0.29 | 0.66 |
| Temporal_Mid_R | -1.00 | 0.32 | 0.77 | -0.21 | 0.83 | 0.96 | -1.56 | 0.12 | 0.57 |
| Temporal_Pole_Mid_L | 0.46 | 0.65 | 0.88 | -0.42 | 0.68 | 0.90 | 0.33 | 0.74 | 0.87 |
| Temporal_Pole_Mid_R | 1.46 | 0.14 | 0.57 | -1.25 | 0.21 | 0.65 | 1.52 | 0.13 | 0.57 |
| Temporal_Inf_L | -0.64 | 0.52 | 0.88 | 0.06 | 0.96 | 0.97 | -0.96 | 0.34 | 0.71 |
| Temporal_Inf_R | 0.36 | 0.72 | 0.89 | -1.42 | 0.16 | 0.51 | -0.31 | 0.75 | 0.87 |
| Cerebelum_Crus1_L | -0.47 | 0.64 | 0.88 | 0.44 | 0.66 | 0.90 | -0.49 | 0.63 | 0.87 |
| Cerebelum_Crus1_R | 0.40 | 0.69 | 0.88 | -0.62 | 0.54 | 0.84 | 0.07 | 0.94 | 0.95 |
| Cerebelum_Crus | 0.87 | 0.39 | 0.80 | -0.26 | 0.80 | 0.95 | 1.22 | 0.22 | 0.58 |

| | | | | | | | | | |
|---|---|---|---|---|---|---|---|---|---|
| 2_L | | | | | | | | | |
| Cerebelum_Crus 2_R | 0.68 | 0.50 | 0.88 | -0.54 | 0.59 | 0.88 | 0.82 | 0.41 | 0.76 |
| Cerebelum_3_L | -0.95 | 0.34 | 0.79 | 0.81 | 0.42 | 0.74 | -0.72 | 0.47 | 0.78 |
| Cerebelum_3_R | -0.59 | 0.55 | 0.88 | 1.00 | 0.32 | 0.68 | 0.09 | 0.93 | 0.95 |
| Cerebelum_4_5_L | 1.54 | 0.12 | 0.55 | -1.43 | 0.15 | 0.51 | 1.39 | 0.16 | 0.57 |
| Cerebelum_4_5_R | 1.71 | 0.09 | 0.44 | -2.02 | 0.04 | 0.40 | 1.24 | 0.22 | 0.58 |
| Cerebelum_6_L | -0.39 | 0.70 | 0.88 | -0.02 | 0.98 | 0.98 | -0.90 | 0.37 | 0.71 |
| Cerebelum_6_R | -0.54 | 0.59 | 0.88 | 0.85 | 0.40 | 0.73 | -0.34 | 0.73 | 0.87 |
| Cerebelum_7b_L | 1.98 | 0.05 | 0.35 | -1.72 | 0.09 | 0.42 | 2.07 | 0.04 | 0.29 |
| Cerebelum_7b_R | 0.71 | 0.48 | 0.87 | 0.09 | 0.93 | 0.97 | 1.30 | 0.20 | 0.58 |
| Cerebelum_8_L | -0.60 | 0.55 | 0.88 | 0.09 | 0.93 | 0.97 | -1.07 | 0.29 | 0.66 |
| Cerebelum_8_R | 0.92 | 0.36 | 0.79 | -1.10 | 0.27 | 0.67 | 0.43 | 0.67 | 0.87 |
| Cerebelum_9_L | -0.07 | 0.94 | 0.97 | -0.08 | 0.94 | 0.97 | 0.21 | 0.84 | 0.92 |
| Cerebelum_9_R | 1.41 | 0.16 | 0.60 | -1.22 | 0.22 | 0.67 | 1.51 | 0.13 | 0.57 |
| Cerebelum_10_L | -0.45 | 0.65 | 0.88 | 0.10 | 0.92 | 0.97 | -0.66 | 0.51 | 0.80 |
| Cerebelum_10_R | 0.27 | 0.78 | 0.92 | -0.24 | 0.81 | 0.95 | 0.43 | 0.66 | 0.87 |
| Vermis_1_2 | -0.92 | 0.36 | 0.79 | 0.26 | 0.80 | 0.95 | -1.19 | 0.23 | 0.59 |
| Vermis_3 | 2.98 | $3.14 \times 10^{-3}$ | 0.07 | -2.80 | $5.62 \times 10^{-3}$ | 0.15 | 2.60 | 0.01 | 0.15 |
| Vermis_4_5 | 1.17 | 0.24 | 0.74 | -1.19 | 0.24 | 0.67 | 1.08 | 0.28 | 0.66 |
| Vermis_6 | 2.04 | 0.04 | 0.35 | -2.62 | $9.63 \times 10^{-3}$ | 0.16 | 1.26 | 0.21 | 0.58 |

| | | | | | | | | | |
|---|---|---|---|---|---|---|---|---|---|
| Vermis_7 | -0.86 | 0.39 | 0.80 | 0.10 | 0.92 | 0.97 | -1.14 | 0.26 | 0.62 |
| Vermis_8 | -0.27 | 0.78 | 0.92 | 0.05 | 0.96 | 0.97 | -0.29 | 0.77 | 0.88 |
| Vermis_9 | -1.10 | 0.27 | 0.76 | 0.63 | 0.53 | 0.84 | -0.84 | 0.40 | 0.76 |
| Vermis_10 | -0.87 | 0.39 | 0.80 | -0.44 | 0.66 | 0.90 | -1.65 | 0.10 | 0.53 |

**Table S16. Samples from the REST-meta-MDD project: consortium sites, sample size, and data acquisition parameters**

| Site | Center | N | | Scanner | TR (ms) | TE (ms) | FA (°) | FOV (mm²) | Resolution (mm²) | Slices | Thickness (mm) | Gap (mm) | Time points |
|---|---|---|---|---|---|---|---|---|---|---|---|---|---|
| | | MDD | NC | | | | | | | | | | |
| 1 | PKU | 73 | 73 | Siemens Tim Trio 3T | 2000 | 30 | 90 | 210×210 | 3.28×3.28 | 30 | 4 | 0.8 | 210 |
| 2 | SCU | 16 | 14 | PHILIPS Achieva 3T | 2000 | 30 | 90 | 240×240 | 1.67×1.67 | 37 | 4 | 0 | 200 |
| 4 | CSU1 | 18 | 23 | Siemens Skyra 3T | 2500 | 25 | 90 | 240×240 | 3.75×3.75 | 39 | 3.5 | 0 | 200 |
| 7 | ZJU | 35 | 37 | GE discovery MR750 | 2000 | 30 | 90 | 220×220 | 2.29×2.29 | 37 | 3.2 | 0 | 184 |
| 8 | CMU | 39 | 48 | GE | 200 | 30 | 9 | 240×2 | 3.75×3. | 35 | 3 | 0 | 200 |

| | | | | | | | | | | | | | |
|---|---|---|---|---|---|---|---|---|---|---|---|---|---|
| | | | | Signa 3T | 0 | | 0 | 40 | 75 | | | | |
| 9 | JNU | 48 | 48 | GE Discovery MR750 | 2000 | 25 | 90 | 240×240 | 3.75×3.75 | 35 | 3 | 1 | 200 |
| 10 | SXMU | 45 | 26 | Siemens Tim Trio 3T | 2000 | 30 | 90 | 240×240 | 3.75×3.75 | 32 | 3 | 1.52 | 212 |
| 11 | CQMU1 | 20 | 17 | GE Signa 3T | 2000 | 30 | 90 | 240×240 | 3.75×3.75 | 33 | 5 | 0 | 200 |
| 13 | XJTU | 20 | 16 | GE Excite 1.5T | 2500 | 35 | 90 | 256×256 | 4.00×4.00 | 35 | 4 | 0 | 150 |
| 14 | CSU2 | 61 | 32 | Siemens Tim Trio 3T | 2000 | 25 | 90 | 240×240 | 3.75×3.75 | 39 | 3.5 | 0 | 200 |
| 15 | SEU | 30 | 37 | Siemens Verio 3T | 2000 | 25 | 90 | 240×240 | 3.75×3.75 | 36 | 4 | 0 | 240 |
| 17 | CQMU2 | 41 | 41 | GE Signa 3T | 2000 | 40 | 90 | 240×240 | 3.75×3.75 | 33 | 4 | 0 | 240 |
| 19 | AHMU | 18 | 31 | GE | 200 | 22. | 3 | 220×2 | 3.44×3. | 33 | 4 | 0.6 | 240 |

| | | | | | | | | | | | | | |
|---|---|---|---|---|---|---|---|---|---|---|---|---|---|
| | | | | Signa 3T | 0 | 5 | 0 | 20 | 44 | | | | |
| 20 | SWU | 245 | 225 | Siemens Tim Trio 3T | 2000 | 30 | 90 | 220×220 | 3.44×3.44 | 32 | 3 | 1 | 242 |
| 21 | CAMU | 79 | 65 | Siemens Tim Trio 3T | 2000 | 30 | 90 | 200×200 | 3.12×3.12 | 33 | 3.5 | 0.7 | 240 |
| 22 | CSU3 | 18 | 20 | Philips Gyroscan Achieva 3.0T | 2000 | 30 | 90 | 240×240 | 1.67×1.67 | 36 | 4 | 0 | 250 |
| 23 | SCU | 22 | 23 | Philips Achieva 3.0T TX | 2000 | 30 | 90 | 240×240 | 3.75×3.75 | 38 | 4 | 0 | 240 |

Abbreviation, PKU: National Clinical Research Center for Mental Disorders (Peking University Sixth Hospital) & Key Laboratory of Mental Health, Ministry of Health (Peking University); CSU: The Second Xiangya Hospital of Central South University; ZJU: Sir Run Run Shaw Hospital, Zhejiang University School of Medicine; CMU: Department of Psychiatry, First Affiliated Hospital, China Medical University; JNU: The First Affiliated Hospital of Jinan University; SXMU: First Hospital of Shanxi Medical University; CQMU: Department of Psychiatry, The First Affiliated Hospital of Chongqing Medical University; XJTU: The First Affiliated Hospital of Xi’an Jiaotong University, Xi’an Central Hospital; SEU: Department of Psychosomatics and Psychiatry, Zhongda Hospital, School of Medicine, Southeast University; AHMU: Anhui Medical University; SWU: Faculty of Psychology, Southwest University; SCU: Mental Health Center, West China Hospital, Sichuan University; SCU: Department of Clinical Psychology, Suzhou Suzhou Psychiatric Hospital, The

Affiliated Guangji Hospital of Soochow University; CAMU: Beijing Anding Hospital, Capital Medical University.
